# The USH3A causative gene clarin1 functions in Müller glia to maintain retinal photoreceptors

**DOI:** 10.1101/2024.02.29.582878

**Authors:** Hannah J. T. Nonarath, Samantha L. Simpson, Tricia L. Slobodianuk, Ross F. Collery, Astra Dinculescu, Brian A. Link

**Affiliations:** Department Cell Biology, Neurobiology and Anatomy, Medical College of Wisconsin, Milwaukee, Wisconsin 53226; Department of Ophthalmology and Vision Sciences, Medical College of Wisconsin, Milwaukee, Wisconsin 53226; Department of Ophthalmology, University of Florida, Gainesville, Florida 32611

## Abstract

Mutations in *CLRN1* cause Usher syndrome type IIIA (USH3A), an autosomal recessive disorder characterized by hearing and vision loss, and often accompanied by vestibular balance issues. The identity of the cell types responsible for the pathology and mechanisms leading to vision loss in USH3A remains elusive. To address this, we employed CRISPR/Cas9 technology to delete a large region in the coding and untranslated (UTR) region of zebrafish *clrn1*. Retina of *clrn1* mutant larvae exhibited sensitivity to cell stress, along with age-dependent loss of function and degeneration in the photoreceptor layer. Investigation revealed disorganization in the outer retina in *clrn1* mutants, including actin-based structures of the Müller glia and photoreceptor cells. To assess cell-specific contributions to USH3A pathology, we specifically re-expressed *clrn1* in either Müller glia or photoreceptor cells. Müller glia re-expression of *clrn1* prevented the elevated cell death observed in larval *clrn1* mutant zebrafish exposed to high-intensity light. Notably, the degree of phenotypic rescue correlated with the level of Clrn1 re-expression. Surprisingly, high levels of Clrn1 expression enhanced cell death in both wild-type and *clrn1* mutant animals. However, rod- or cone-specific Clrn1 re-expression did not rescue the extent of cell death. Taken together, our findings underscore three crucial insights. First, *clrn1* mutant zebrafish exhibit key pathological features of USH3A; second, Clrn1 within Müller glia plays a pivotal role in photoreceptor maintenance, with its expression requiring controlled regulation; third, the reliance of photoreceptors on Müller glia suggests a structural support mechanism, possibly through direct interactions between Müller glia and photoreceptors mediated in part by Clrn1 protein.

**AUTHOR SUMMARY:** Mutations in USH-associated genes profoundly impact patients, affecting auditory, visual, and vestibular function. While the basis of inner ear defects is reasonably well understood for USH and auditory devices can improve hearing, the mechanisms underlying photoreceptor loss are unknown, and there are no approved treatments for vision deficits. In USH3A, the affected gene, *clarin1* (*clrn1*), is predominantly expressed in Müller glia. The role of Müller glia in maintaining photoreceptor health and contributions to USH3 pathology is understudied, in part as *Clrn1* mutant mice - the traditional experimental model used to study retinal diseases - do not phenocopy the photoreceptor loss of USH3 patients. In the present study, we developed a zebrafish model of USH3A that displays many features of the human disease. Our research shows that the loss of Clrn1 affects actin-based structures of the outer retina, including those of photoreceptor cells and Müller glia. Importantly, we demonstrate that the expression of Clrn1 in Müller glia, but not rods and cones, alleviated light-induced damage in *clrn1* mutant zebrafish. We also highlight that the dosage of Clrn1 in Müller glia is critical for maintaining proper photoreceptor function. These findings demonstrate the key contribution of Müller glia to USH pathology and can guide strategies for gene-replacement therapies.

## INTRODUCTION

Usher syndrome (USH) is an autosomal recessive disorder characterized by the loss of both hearing and vision. This syndrome accounts for more than half of all hereditary cases of deaf–blindness in the United States (1–4). USH can be subdivided into three clinical classifications (USH1, USH2, USH3) based on the onset, severity, and progression of hearing loss and vestibular abnormalities. USH is further subtyped depending on the affected gene. For all clinical classifications, hearing loss is the primary identifying symptom. Vision loss presents as retinitis pigmentosa, beginning with the loss of night vision followed by a gradual narrowing of the visual field and blindness.

Although all mouse models of USH display early-onset hearing loss, most fail to present with degenerative retinal phenotypes. This represents a major challenge, hampering the investigation of mechanisms leading to vision loss (5–7). Potential reasons for the absence of retinal degeneration in murine models include marked interspecies differences in photoreceptor ultrastructure, rod/cone ratio, density and distribution, and a potentially distinct subcellular distribution of some of the USH proteins (8). Specifically, the mouse retina is rod dominant and the photoreceptors lack well-defined calyceal processes, which have previously been described as being abnormal or absent (8,9). The calyceal processes are long microvilli-like actin rich structures resembling hair cell stereocilia, that emerge from the apical region of the inner segment (IS) and envelop the base of the outer segment (OS) in rods and cones. This collar-like structure of the calyceal processes is present in zebrafish, frogs, pigs and primates, including humans, but is absent or vestigial in mouse photoreceptors (8,10,11). Within the human and non-human primate retina, USH1 proteins were reported to localize to the calyceal processes and the inner–outer segment interface (8,12). The USH1 protein Harmonin was also localized at the tips of the Müller glia apical microvilli and the adherens junctions between Müller glia and photoreceptor cells forming the outer limiting membrane in the human retina (13). Harmonin localization is consistent with a previous study performed in zebrafish retina (14). While localization of USH proteins is an important first step in understanding their functions, genetic mutation studies in models capable of replicating the photoreceptor degeneration are critical to understand the underlying mechanisms of USH pathology. Notably, zebrafish loss-of-function models of USH1 and USH2 genes orthologs have been instrumental in characterizing their roles in photoreceptor structure and maintenance (14–18). In contrast, virtually nothing is known about the mechanisms of retinal degeneration associated with USH3.

Mutations in *CLRN1* gene (clarin1, *USH3A*) encoding a four-transmembrane domain protein (CLRN1), are the leading cause of USH3, resulting in progressive loss of hearing and retinitis pigmentosa in humans, with variable vestibular dysfunction (19–22). As with other USH-associated proteins, understanding the role of CLRN1 in the retina has been hindered by the lack of a retinal degenerative phenotype in murine models and discrepancies in its reported cellular expression (23). Specifically, both the *Clrn1* KO and N48K knock-in mouse models of USH3A present with an early-onset hearing loss and are profoundly deaf by postnatal day P30 (7,24). However, both models lack a retinal degeneration phenotype (7,24). Our understanding of CLRN1 function primarily comes from cell-culture studies and research of the inner ear in mouse and zebrafish models (7,25–27). Cell culture experiments have shown that CLRN1 is trafficked to the plasma membrane and may potentially function as a regulator of the actin organization (27). In mouse and zebrafish models of USH3A (*clrn1*), the mutant animals presented with disorganized stereocilia bundles, which are sensory organelles composed of actin-rich protrusions on the apical surface of the auditory and vestibular sensory hair cells (7,28–31).

Furthermore, the loss of clarin1 also led to the disorganization of the synaptic actin network in the inner hair cells in a mouse model of USH3 (31). By 7 days post fertilization (dpf), previously established USH3A zebrafish mutants presented with hearing deficits and splayed hair cells (28). These mutant animals did not survive into adulthood and there were no changes in retinal function in larvae as measured by electroretinogram (ERG) analysis (28). Within the inner ear, the disorganization of hair cell stereocilia prevented the proper transduction of mechanical force from sound, head movement, or gravity into electrical signals. These findings suggest that CLRN1 regulates the formation and maintenance of properly shaped hair bundles in both the outer and inner hair cells of the inner ear (26). Whether CLRN1 performs a similar role in maintaining the integrity of actin rich structures in the retina is unknown.

Published studies indicate Clrn1 transcripts are abundantly enriched in Müller glia, as compared to other cell types. In the early postnatal mouse retina, mRNA in-situ hybridization analysis revealed that *Clrn1* expression was restricted to the INL and co-localized with Müller glia cell-specific markers (7,32). In a cross-species study, in-situ-hybridization and single-cell RNAseq analysis revealed that *Clrn1* transcripts in the mouse and human adult retina were concentrated to the INL and specifically enriched in Müller glia (32). Other studies based on scRNAseq analysis have also reported that *Clrn1* mRNA is enriched in retinal progenitors and Müller glia across several species, including human, non-human primate, and zebrafish (33–35) . Transcriptomic studies suggest that several USH1 protein, including HARMONIN (USH1C), CDH23 (USH1D), and SANS (USH1G) are present in Müller glia (33). Additionally, *in vitro* binding studies suggest that CLRN1 and HARMONIN may directly interact (31). Uncovering the roles of Müller glia-expressed USH proteins in the retina will shed light not only on mechanisms of USH pathology, but also advance our understanding of the relationship between Müller glia and photoreceptors.

In this study, we developed an USH3A zebrafish model by deleting the *clarin1* locus. Unlike the previously reported USH3A zebrafish models, homozygous mutants developed for this study are viable and survive into adulthood, enabling the identification of a photoreceptor degeneration phenotype and a detailed characterization of the importance of Müller glia in USH3 ocular pathology.

## RESULTS

### Development of *clrn1* mutant zebrafish

To develop an USH3A model in zebrafish, 85% of the coding sequence for *clrn1* was deleted through a CRISPR-Cas9 strategy. Specifically, we used one CRISPR gRNA that targeted a region of exon 1 and another CRISPR gRNA designed to target the 3’ UTR of *clrn1* (Figure 1a). To confirm the deletion and establish a genotyping protocol, primers flanking the cut sites were designed to discriminate the genomic deletion (Supplemental Figure 1a). Additionally, Sanger sequencing (Supplemental Figure 1b) and RNAscope for *clrn1* (Figure 1c) were applied to confirm the *clrn1* deletion. Sanger sequencing demonstrated the elimination of the genomic region between the targeted CRISPR cut sites, while RNAscope analysis substantiated the complete loss of *clrn1* transcripts (Supplemental Figure 1b, Figure 1c). Furthermore, RNAscope showed that in zebrafish, similar to other species, *clrn1* transcripts concentrate in the inner nuclear layer of the retina. Lastly, to confirm the loss of Clrn1 function, we evaluated previously established hair cell phenotypes of the inner ear associated with USH3 models (7,28). At 7 dpf, the hair cell organization was grossly altered, with *clrn1^-/-^*zebrafish larvae presenting with splayed stereocilia (Figure 1d). In some animals, pyknotic nuclei were also observed in the hair cells of the anterior macula at the time point investigated (Figure 1d). However, in *clrn1^-/-^* larvae no severe vestibular phenotype was observed, such as the circular swimming pattern reported in the *myo7aa* and *pcdh15a* mutant zebrafish, which model USH1B and USH1F, respectively (36–38).

**Figure 1:**
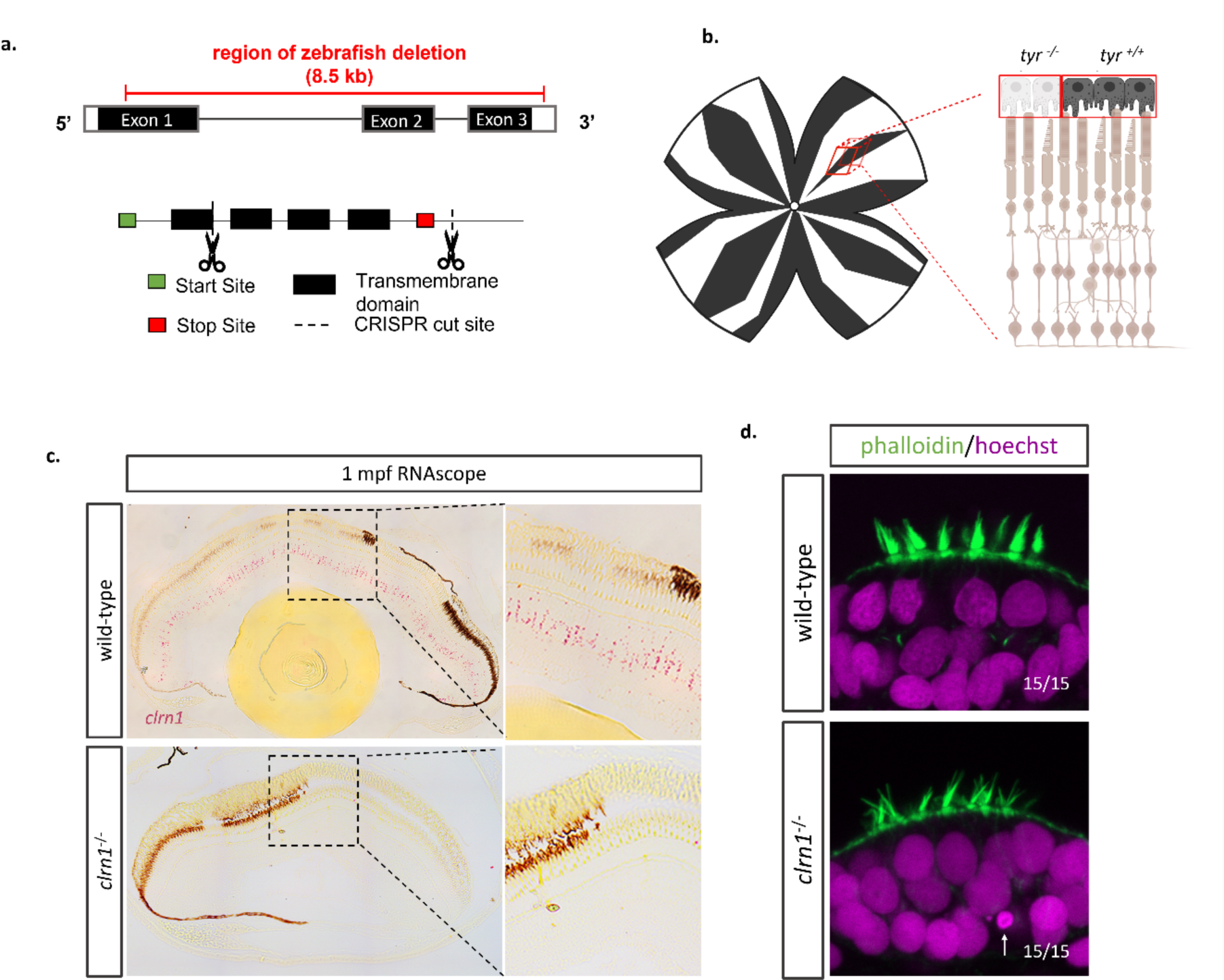
Development of USH3A zebrafish model. A *clrn1* mutant zebrafish was generated using CRISPR/Cas9 technology. Pairs of CRISPR gRNAs were designed to target Exon 1 and the 3’ UTR of *clrn1*. (a) Depiction of CRISPR cut sites in relation to *clrn1* zebrafish exons and protein functional domains. (b) Method to mosaically deplete pigmentation from the retinal pigment epithelium to allow for RNAscope probe detection. (c) Detection of *clrn1* transcripts (bright red dots) in the inner retina of 1 mpf wild-type zebrafish using the highly sensitive RNAscope in-situ hybridization assay. (d) Whole-mount phalloidin staining on 7dfp wild-type and *clrn1^-/-^* larvae to assess hair cell structure. Arrow indicates pycnotic nuclei.

### *Clrn1^-/-^* adult zebrafish show progressive disruption of the cone photoreceptor mosaic

Several established zebrafish models of USH do not survive into adulthood, including previously generated *clrn1* mutant zebrafish, preventing the study of disease progression and pathophysiology (28). However, the *clrn1^-/-^* zebrafish developed for this study do survive into adulthood, providing an opportunity to characterize aged retinal phenotypes. To assess whether mutant retinas show the progressive photoreceptor disorganization and degeneration that is observed in USH3A patients, we utilized optical coherence tomography (OCT), which can provide single cell resolution of photoreceptor cell organization. Retinas of wild-type and *clrn1^-/-^*zebrafish were imaged at 4-, 8-, 12-, and 20-months post fertilization (mpf). From the OCT scans, *en face* images of the ultraviolet-sensitive (UV) cone photoreceptor packing arrangement and spacing regularity were assessed by Voronoi cell area, number of neighbors, and intercell distance regularity (39). Alterations in these measurements are an indication of disrupted photoreceptor structure and/or survival (39). As expected in wild-type zebrafish, a characteristic crystalline lattice of linear columns and rows of UV cone photoreceptors was observed (Figure 2a-d). At 4 and 8 mpf, *clrn1^-/-^* zebrafish also displayed the highly regular photoreceptor arrangement, although at 8 mpf, there were a few discernable gaps in the mosaic (Figure 2e-f). By 12 mpf, the *clrn1^-/-^* zebrafish exhibited more areas where signals generated by the photoreceptor outer segments were absent (Figure 2g-g’). At 20 mpf, an OCT reflective signal from cone outer segments was completely absent in 15% of the *clrn1^-/-^* zebrafish (Figure 2h). Among the *clrn1^-/-^*retinas that still exhibited discrete reflective UV cone outer segments, the organization was significantly disrupted, as indicated by the loss of the hexagonal patterning in the Voronoi overlay (Figure 2h’’). In contrast, smaller changes, as part of normal ageing, occurred to the UV cone mosaic of wild-type animals from 4 to 20 mpf (Figure 2a-d). Collectively, the OCT data suggest age-dependent photoreceptor changes due to loss of Clrn1.

**Figure 2:**
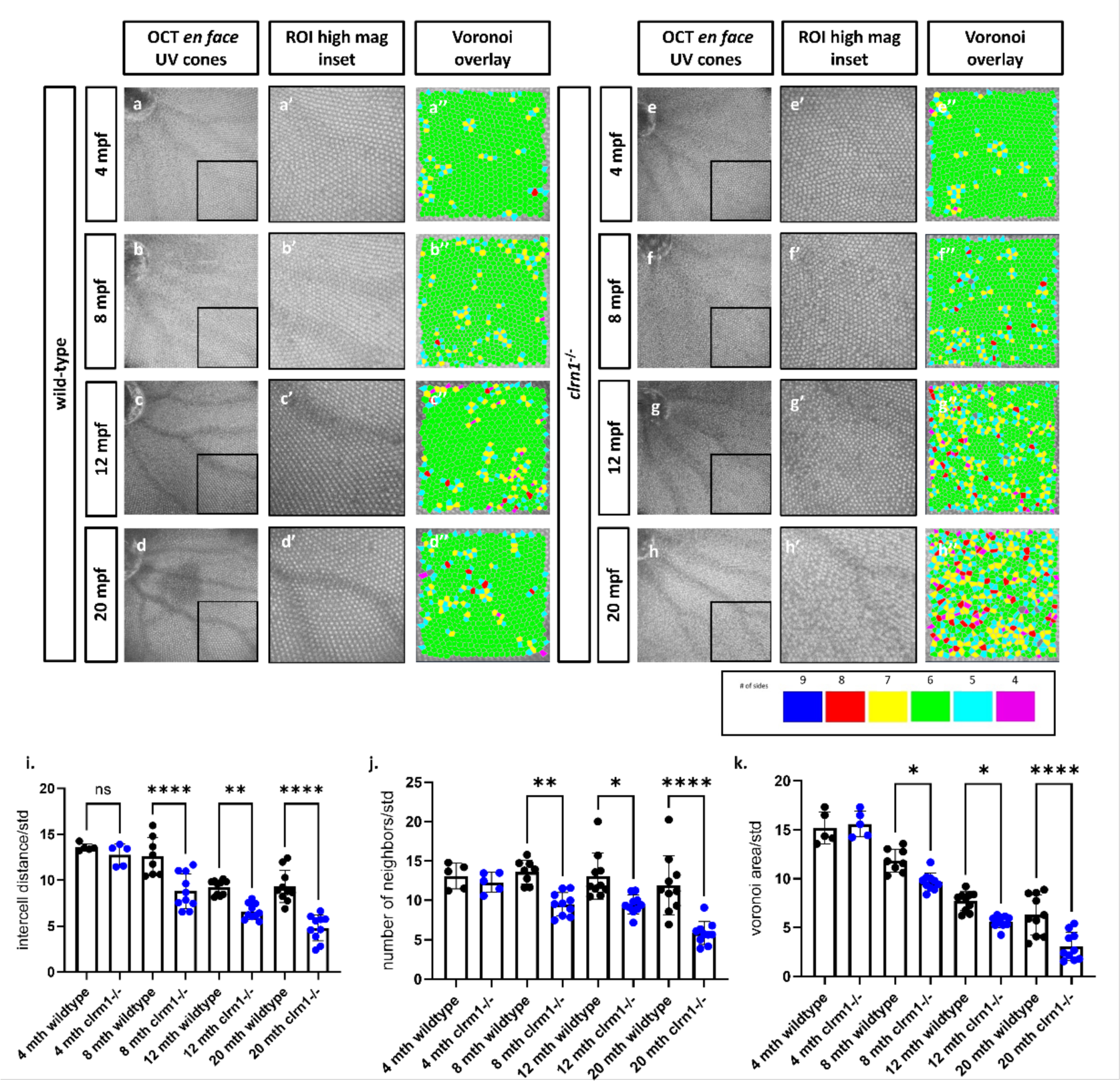
*clrn1^-/-^* zebrafish present with an altered UV cone mosaic beginning at 8 mpf. *En face* images of the UV cone mosaic were segmented from OCT scans and process for the Voronoi overlay in wild-type at (a) 4, (b) 8, (c) 12, and (d) 24 mpf, as well as *clrn1^-/-^* zebrafish at (e) 4, (f) 8, (g) 12, and (h) 24 mpf. (i) Quantification of measures of the UV cone mosaic regularity revealed statistically significant differences in the intercell distance regularity, (j) number of neighbors regularity and, (k) the Voronoi area regularity at 8 mpf in *clrn1^-/-^* zebrafish compared to wild-type. (*p<0.05, **p<0.01, ****p<0.001; Two-way ANOVA)

### Photopic and scotopic ERG responses are abnormal in *clrn1^-/-^* zebrafish

To assess whether the loss of Clrn1 results in retinal dysfunction, we conducted *ex vivo* electroretinography (ERG) on retinas collected from 4- and 12-mpf wild-type or *clrn1^-/-^* zebrafish. Our findings revealed discernible functional differences in both scotopic and photopic responses at these time points, as illustrated in Figure 3. Notably, we observed considerable variability in the scotopic b-wave among *clrn1^-/-^* fish, with 30% of the *clrn1^-/-^* fish displaying amplitudes greater than those observed in the 4 mpf wild-type zebrafish (Supplemental Figure 2a). Additionally, significant reductions in the photopic b-wave were observed in 4 mpf *clrn1^-/-^* zebrafish (Figure 3b). Furthermore, substantial differences were noted in the amplitude of the flicker responses at lower frequencies (Figure 3c and Supplemental Figure 3). Collectively, these findings suggest functional disparities in visual transduction in 4 mpf *clrn1^-/-^* animals.

**Figure 3.**
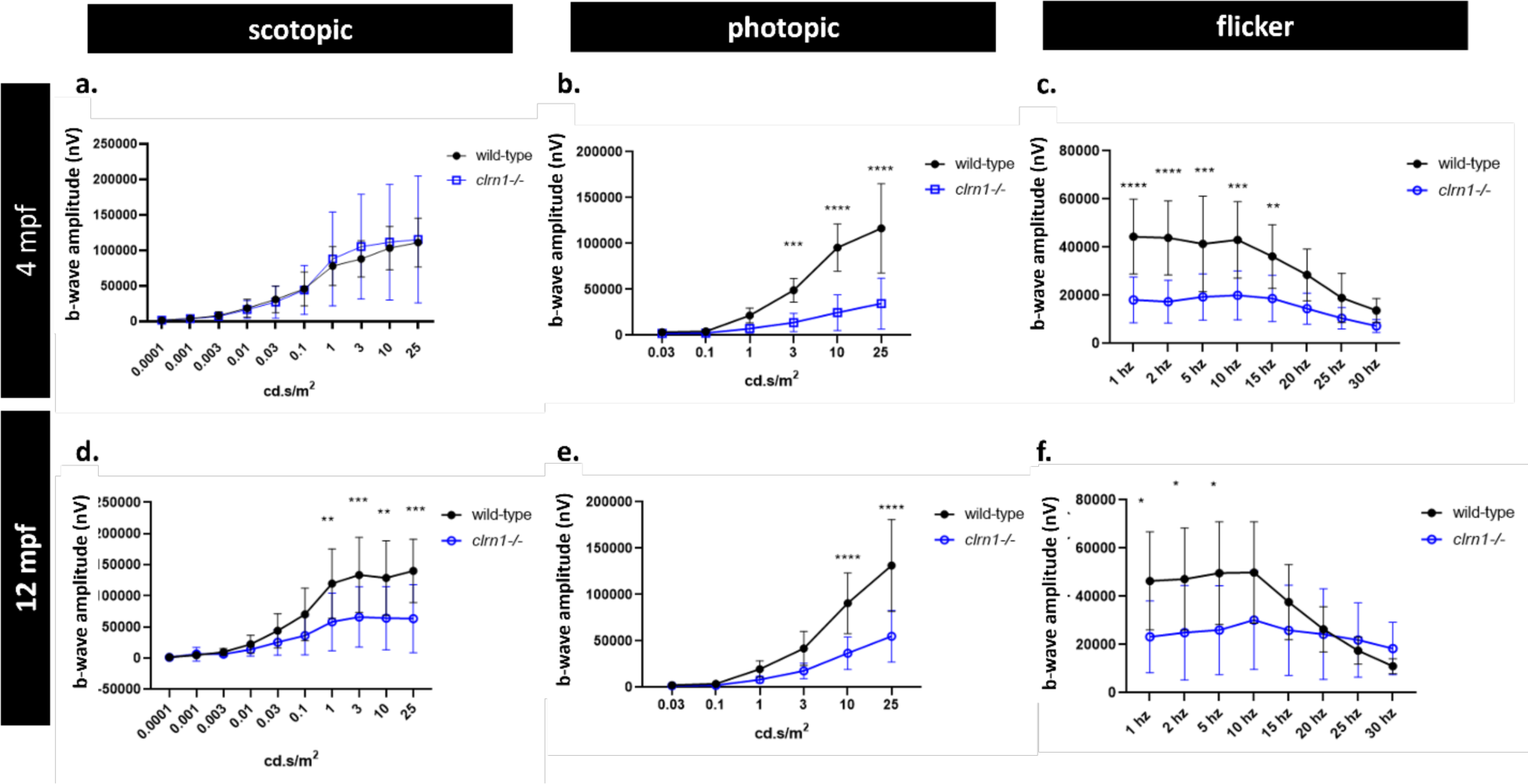
Scotopic and photopic ERG responses are affected by the loss of Clrn1. Quantification of 4 mpf wild-type and *clrn1^-/-^* b-wave amplitudes for ERG scotopic (a) and photopic (b) responses. Quantification of 4 mpf flicker b-wave amplitude for wildtype and *clrn1^-/-^* zebrafish (C). Quantification of 12 mpf wild-type and *clrn1^-/-^* b-wave amplitudes for ERG scotopic (d) and photopic (e) responses. Quantification of 4 mpf flicker bwave amplitude for wildtype and clrn1-/-zebrafish (f). (**p<0.01, ***p<0.005, ****p<0.001; One-way ANOVA). Error bars=SD.

Similar distinctions in scotopic and photopic responses were evident in the 12 mpf *clrn1^-/-^* zebrafish when compared to their age-matched wild-type counterparts (Figure 3d-e). At 12 mpf, the scotopic b-waves were consistently diminished, and less variable (Figure 3d and Supplemental Figure 2c). Furthermore, the significant reduction in the photopic b-wave, coupled with a marked decrease in the photopic flicker response, persisted in the 12 mpf *clrn1^-/-^* zebrafish (Figure 3d-f). At 12 mpf, an irregular patterning of the flicker response was also observed at 25 Hz and 30 Hz, suggesting potential deficits in the recovery of cone photoreceptor function between stimulations (Supplemental Figure 3a).

In addition to amplitude, we evaluated differences in the scotopic and photopic b-wave implicit time. In 4 mpf fish, no significant difference in implicit times between mutants and wild-type siblings was observed (Supplemental Figure 4a and b). At 12 months of age, however, we did observe a significant increase in the photopic b-wave implicit time in *clrn1^-/-^*zebrafish (Supplemental Figure 4d). Overall, our data suggests that the loss of Clrn1 affects both rod and cone functions.

### Slow and progressive photoreceptor loss in *clrn1^-/-^* zebrafish

Differences in the *clrn1^-/-^* zebrafish photoreceptor mosaic, as detected by OCT, may be attributed to cell death or disruption of the photoreceptor outer segments. Similarly, the differences in the ERG responses observed between *clrn1^-/-^* zebrafish and their wild-type siblings could be due to the loss of photoreceptors or the presence of non-functional photoreceptors. Therefore, we performed histological analysis on retinal sections from 4-, 8-, 12-, and 20-mpf wild-type and *clrn1^-/-^* zebrafish to determine if there was a loss of photoreceptor cells. At 20 mpf, we observed obvious thinning of the outer nuclear layer (ONL) in *clrn1^-/-^*zebrafish (Figure 4a), compared to wild-type siblings. We therefore quantified the photoreceptor loss at early stages, and found a slow, but progressive thinning of the ONL starting as early as 4 mpf (Figure 4c). Analysis of rod versus cone loss reveals that the initial decrease in photoreceptor nuclei begins with rod degeneration, and the onset of cone degeneration began by 8 mpf (Figure 4e-f). In addition to the decrease in nuclei within the ONL, there was also a shortening of the rod outer segments, a phenotype frequently observed in models of retinal degeneration (40–42) (Figure 4b and c).

**Figure 4:**
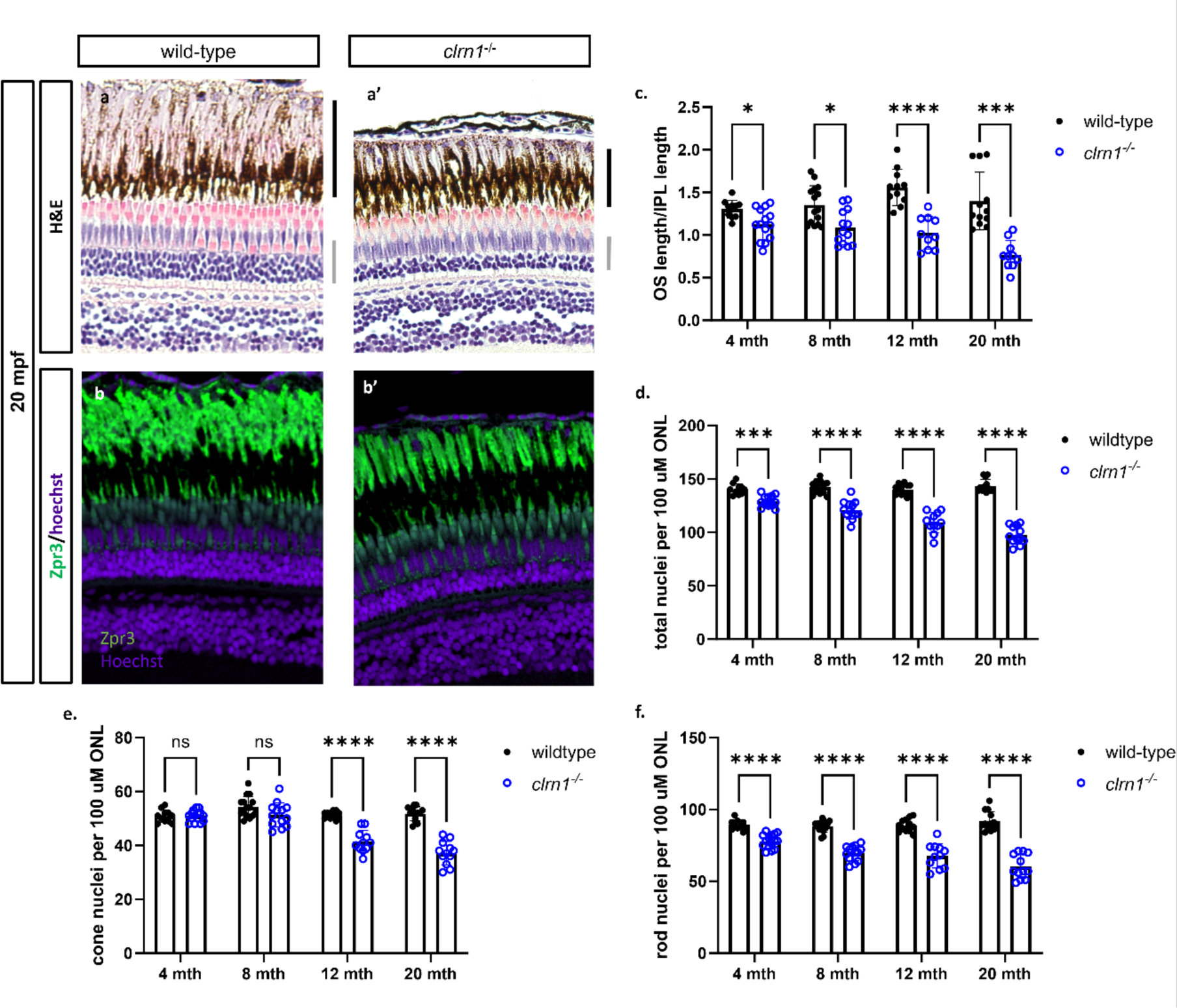
*clrn1^-/-^* zebrafish present with age-dependent shortening of the rod outer segments and thinning of the outer nuclear layer. Hematoxylin and eosin-stained paraffin section from 20 mpf wild-type and *clrn1^-/-^* zebrafish (a). Staining for Zpr3 (green) on paraffin sections from 20 mpf wild-type and *clrn1^-/-^* zebrafish (b). Quantification of rod outer segments (OS) length at 4, 8, 12, and 20 mpf (c) Quantification of total photoreceptors per 100 µm of ONL length at 4, 8, 12, and 20 mpf (d) Analysis of rod (e) and cone (f) photoreceptors in wild-type and *clrn1^-/-^* zebrafish at 4, 8, 12, and 20 mpf. The black bar notes the location of the rod OS, and the grey bar notes the outer nuclear layer. (*p<0.05, **p<0.01, ***p<0.005, ****p<0.001; One-way ANOVA). (n=12-15 per group)

Unlike mammals, zebrafish can regenerate a damaged retina. Therefore, to assess potential cell regeneration we evaluated BrdU incorporation, which is often used to mark regeneration. We detected a small, but significant increase in BrdU-positive cells at 4 mpf (Supplemental Figure 5). At subsequent ages, BrdU incorporation remained moderately elevated (Supplemental Figure 5e and f), despite the overall loss of *clrn1*^-/-^ photoreceptor cells with age. Given the loss of cells in the photoreceptor layer, the observed BrdU-positive cells may represent attempted but inadequate regeneration, and/or dying cells undergoing DNA repair; which has been reported (43–48).

### Enhanced photoreceptor cell death to constant, high-intensity light in *clrn1^-/-^* zebrafish

Previous studies have shown that photoreceptor survival in mutant USH1 and USH2 larvae was compromised in a high-intensity light stress assay (16,17). To determine if similar sensitivities occurred in our USH3A zebrafish model, animals were exposed to constant and high-intensity light to enhance cell stress from 5-7 dpf (Figure 5a). Under standard husbandry conditions, we did not observe histological changes or significant differences in cell death markers in either wild-type or *clrn1* mutant larvae (Figure 5b, 2d-e). Similarly, in the wild-type fish exposed to constant and high-intensity light, we did not observe any histological changes or induction of cell death. Conversely, *clrn1^-/-^* fish exposed to constant and high-intensity light presented with significantly elevated number of pyknotic nuclei in the ONL layer (Figure 5b and 5d), indicating cell death. In addition to the presence of pyknotic nuclei, there was a significant increase in TUNEL-positive nuclei in the ONL, indicative of apoptosis (Figure 5b-c and 5e). We did not observe an increase in the abundance of pyknotic nuclei or TUNEL-positive cells in the other retinal layers of *clrn1^-/-^* fish.

**Figure 5:**
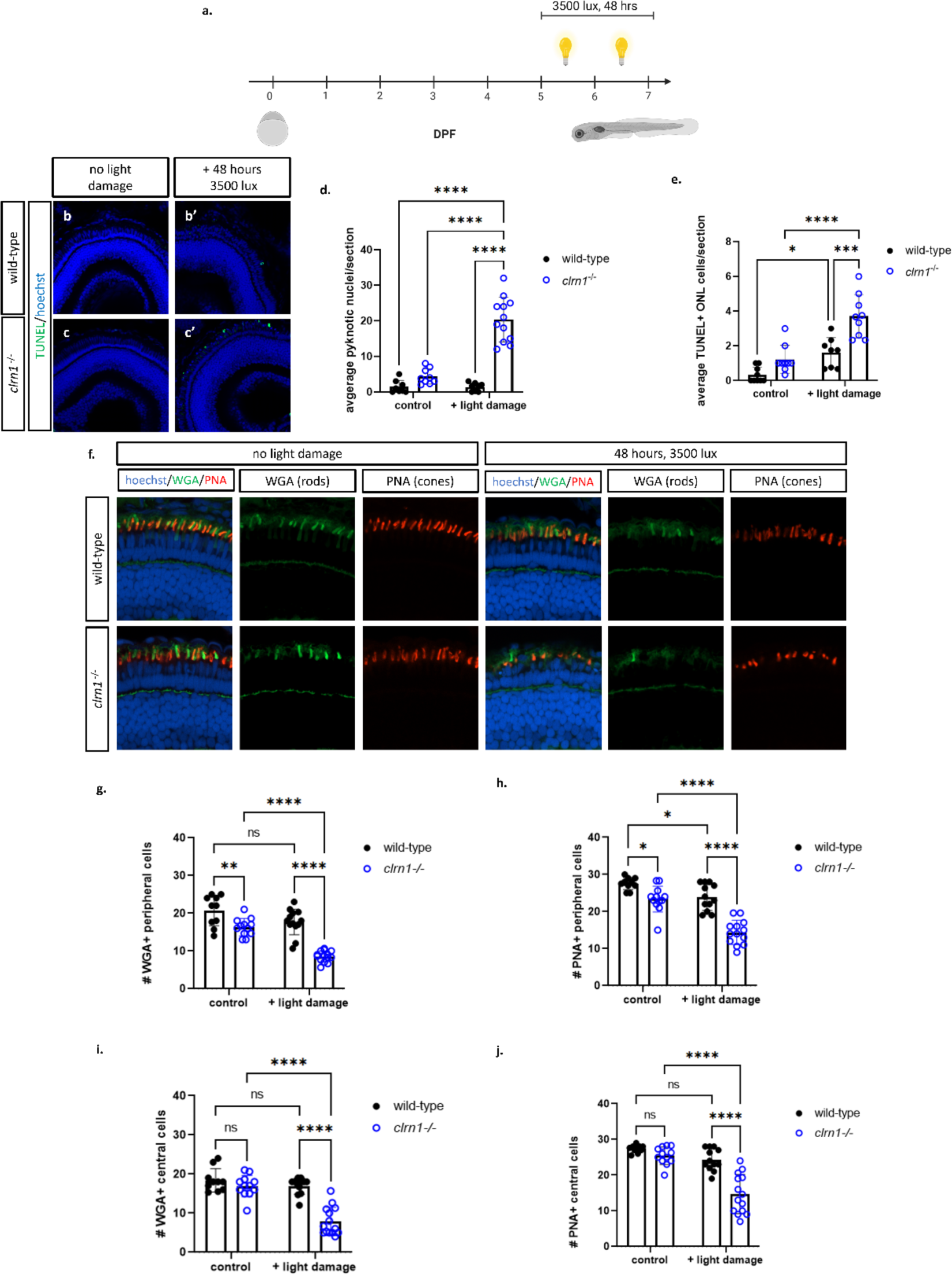
High-intensity light exposure induces rod and cone cell death in *clrn1*^-/-^ zebrafish. Animals were treated with 3500 lux for 48 hours from 5-7 dpf (a). TUNEL staining on transverse cryosectioned retinas from 7 dpf wild-type(b) and *clrn1^-/-^* (c) zebrafish revealed a significant increase in the average number of pyknotic nuclei (d) and TUNEL (e) positive cells per section in the ONL of *clrn1^-/-^* zebrafish. There was no significant increase in TUNEL positive cells detection in the INL (f). Retinal sections from 7dpf control or light-damaged wild-type and *clrn1^-/-^* zebrafish stained with wheat germ agglutinin (WGA, green) and peanut agglutinin (PNA, red) to assess for the loss of rods and cones, respectively (g). Quantification of WGA (h) and PNA (i) positive outer segments along a 100 µm region of the peripheral region revealed a significant decrease in rod and cone outersegments in control *clrn1^-/-^* zebrafish, which was further decreased in light-damaged *clrn1^-/-^* zebrafish. Quantification of WGA (j) and PNA (k) positive outer segments along a 100 µm section of the central region revealed a significant decrease in rod and cone outer segments in light-damaged *clrn1^-/-^* zebrafish. (*p<0.05, **p<0.01, ****p<0.001; Two-way ANOVA). Error bars=SD. Scale Bar: 100 µm (n=10-15 per group)

In addition to elevated cell death, we also evaluated potential functional changes following light damage. Using an optomotor response (OMR) assay, we tested larvae on their ability to detect direction changes. Unexpectedly, we observed functional deficits in the *clrn1^-/-^* zebrafish housed under standard husbandry conditions compared to wild-type animals (Supplemental Figure 6). The failure of mutant larvae to respond to OMR stimulus under control lighting conditions suggests baseline visual impairment.

As evident by ERG results, both rods and cones were affected by the loss of Clrn1. To discern if cell death in the ONL, following light damage, occurred in rods or cones, we stained with fluorescent wheat germ agglutinin (WGA) and peanut agglutinin (PNA), which labels the rod and cone outer segments, respectively. In light-damaged *clrn1^-/-^* zebrafish, there was a significant decrease in WGA and PNA-positive outer segments in the peripheral and central retina of *clrn1^-/-^* zebrafish, compared to that of wild-type siblings (Figure 5f-j). Specifically, we noted a small, but significant decrease in the number of rod and cone outer segments in the peripheral retina of *clrn1^-/-^* 7 dpf larvae in control lighting conditions, which was exacerbated with high-intensity light stress (Figure 5g-h). Collectively, data indicates that at the time point investigated, the rod and cone photoreceptors in *clrn1^-/-^* zebrafish show signs of dysfunction and are sensitized to the cellular stress induced by elevated light exposure.

### Loss of Clrn1 disrupts actin-rich structures of the outer retina

Thus far, results highlight that the loss of Clrn1 in zebrafish induces a slow, progressive photoreceptor dysfunction and degeneration along with sensitization to light-mediated stress. Therefore, we focused on identifying retinal changes associated with the loss of Clrn1. Previous in vitro and in vivo studies suggest a possible role of CLRN1 in the regulation of the F-actin cytoskeleton (27,29–31). To explore the function of Clrn1 in the retina, we first assessed the structure of the ribbon synapses and actin-rich structures of the photoreceptors in wild-type and *clrn1^-/-^* zebrafish. In 7 dpf larvae, we did not find a substantial loss of synapses in the photoreceptor layer, as indicated by WGA staining, which stains pre- and post-synaptic membranes. (Supplemental Figure 7a and b). Ultra-structurally, there were no notable defects in the development of the rod (Supplemental Figure 7c and d) or cone (Supplemental Figure 7e and f) ribbon synapses. Similarly, in the adult retina, we observed comparable *ribeye* staining and appropriate localization of SV2 at the ribbon synapses, further indicating that the ribbon synapses are properly developed in the absence of Clrn1 (Supplemental Figure 7g-i). In summary, the photoreceptor synapses appeared normal in *clrn1*^-/-^ zebrafish.

We next assessed for potential changes in actin structures of the outer retina. Following phalloidin staining, we observed altered actin organization in the photoreceptor layer of *clrn1^-/-^* zebrafish compared to wild-type siblings (Figure 6b). In wild-type fish, actin staining ran perpendicular at the connecting cilium and parallel to the outer segments encompassing the photoreceptor in an actin-rich ring (Figure 6a). In *clrn1*^-/-^ zebrafish larvae, actin staining was disorganized along the outer segment and at the IS/OS junction (Figure 6b). While actin bundles running parallel to the outer segment were evident, the ring structure encompassing the outer segment appeared disorganized.

**Figure 6:**
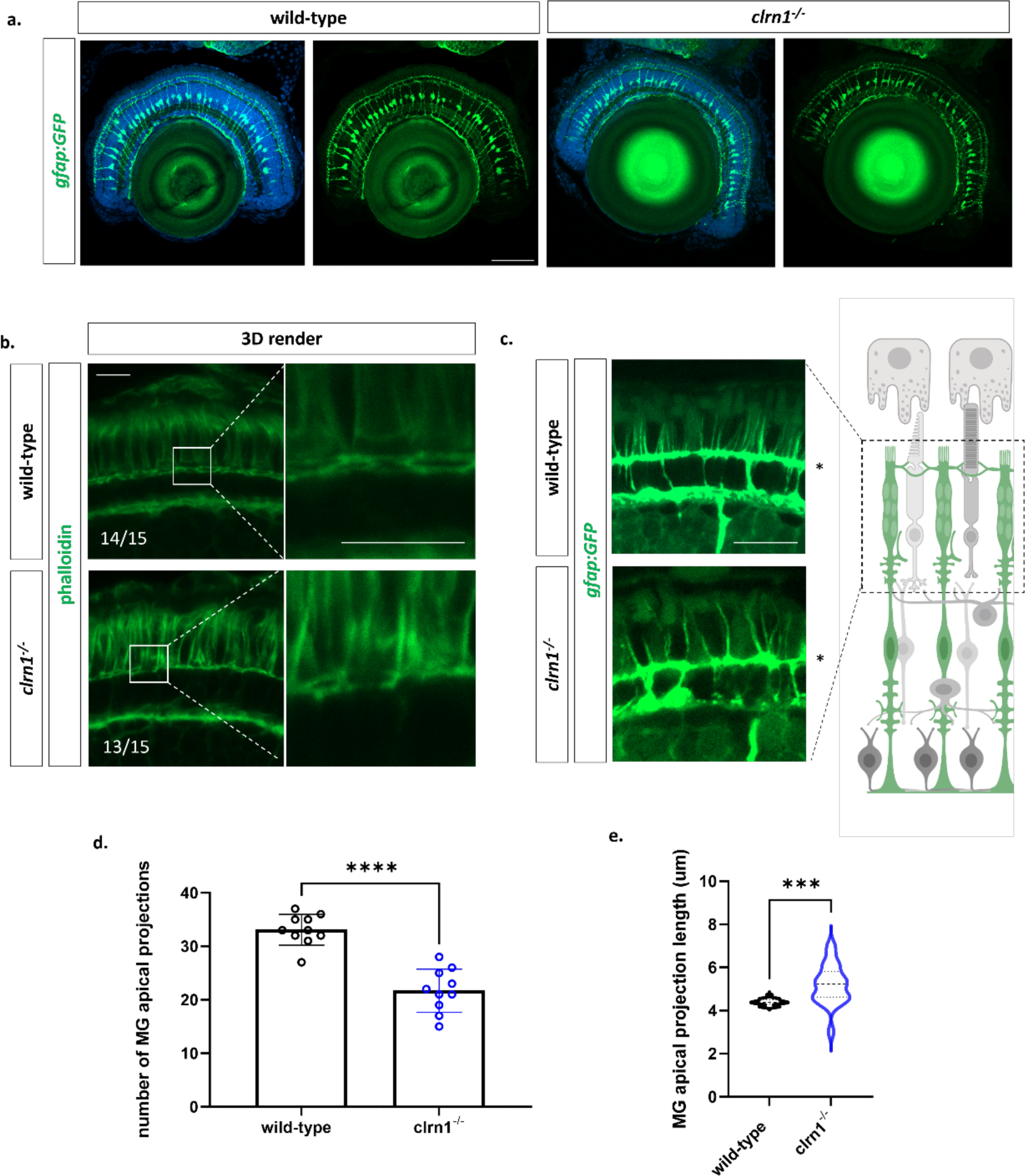
Actin is disorganized in the outer retina of *clrn1^-/-^* zebrafish. Transverse sections from 7 dpf wild-type and *clrn1^-/-^* Tg(*gfap:gfp*) zebrafish (a) highlighting Müller glia expansion from the inner retina to the ONL (a). Phalloidin (green) staining performed on 7 dpf wild-type and *clrn1^-/-^* zebrafish transverse retinal cryosections with high-mag of 3D rendered phalloidin staining for ROI reveals disorganization of actin in the ONL (b) Analysis of MG apical projections within the ONL of wild-type and *clrn1^-/-^*zebrafish shows disorganized MG apical microvilli (c). Quantification of total MG apical microvilli within a 40 µm^2^ region of the central retina of wildtype and *clrn1^-/-^* zebrafish (d). Violin plot of Müller glia apical microvilli length beyond the OLM (*) for wild-type and *clrn1^-/-^*7 dpf zebrafish. (***p<0.005,****p<0.001; Students T-test) Scale bars: 10 µm

On account of changes in actin staining and enriched *clrn1* expression in Müller glia (25,49), we explored in more detail Müller glia structure in 7 dpf larvae. We used a transgenic Müller glia reporter line, which expresses GFP downstream of the glial fibrillary acidic protein (*gfap*) promoter (*Tg(gfap:GFP*). Comparing *clrn1^-/-^* and wild-type zebrafish, we did not observe obvious differences in Müller glia numbers or structure at this age (Figure 8a). Specifically, we observed proper extension of Müller glia processes in both wild-type and *clrn1^-/-^* fish from the INL to the photoreceptor layer, with cell bodies located in the second row of nuclei in the INL (50).

While the abundance and gross morphology of Müller glia were normal in *clrn1^-/-^* mutant zebrafish, the apical microvillar projections were reduced and disorganized. In wild-type larvae, the Müller glia apical microvilli extended beyond the outer limiting membrane (OLM), projecting in a parallel fashion along the photoreceptor outer segments (Figure 6c). These structures were also observed in *clrn1^-/-^* zebrafish but were reduced in number (Fig 6c-d). They also appeared more variable in length, and were less linear in appearance, reminiscent of the splayed stereocilia phenotype of inner ear hair cells (Figure 1d, 6c-d, and 7b).

A common phenotype of several USH1 zebrafish mutants is disrupted photoreceptor calyceal processes (15,51). To investigate the cone calyceal processes of *clrn1*^-/-^ mutants in more detail, we developed a transgenic line, *Tg(gnat2:Lifeact-mCherry*), that expresses Lifeact, an F-actin binding protein (52), under the cone-specific *gnat2* promoter. Transverse cryosections showed enriched staining of Lifeact-mCherry within the calyceal processes, as well as at the adherens-type junctions of the OLM (Fig 7a). Calyceal processes of *clrn1^-/-^* mutants were disorganized and showed a spectrum of phenotypes. The majority of mutant retinas showed elongated, moderately splayed calyceal processes that were reduced in number compared to those of wild-type siblings (Fig 7b). Others were more severely splayed and contained Lifeact-mCherry puncta (Fig 7c). Lifeact-mCherry puncta were also noted at the OLM (Figure 7c’’ and g’’). Given the similarity of phenotypes between mutant Müller glia apical microvilli and the photoreceptor calyceal processes, we combined the *gnat2:Lifeact-mCherry* and *gfap*:*GFP* transgenes to investigate potential associations between these actin-rich structures. We found that in wild-type retina the apical projections of Müller glia were in close association with cone calyceal processes (Fig 7a merged). However, this association was often disrupted in *clrn1^-/-^* retina (Fig7 b,c merged).

**Figure 7:**
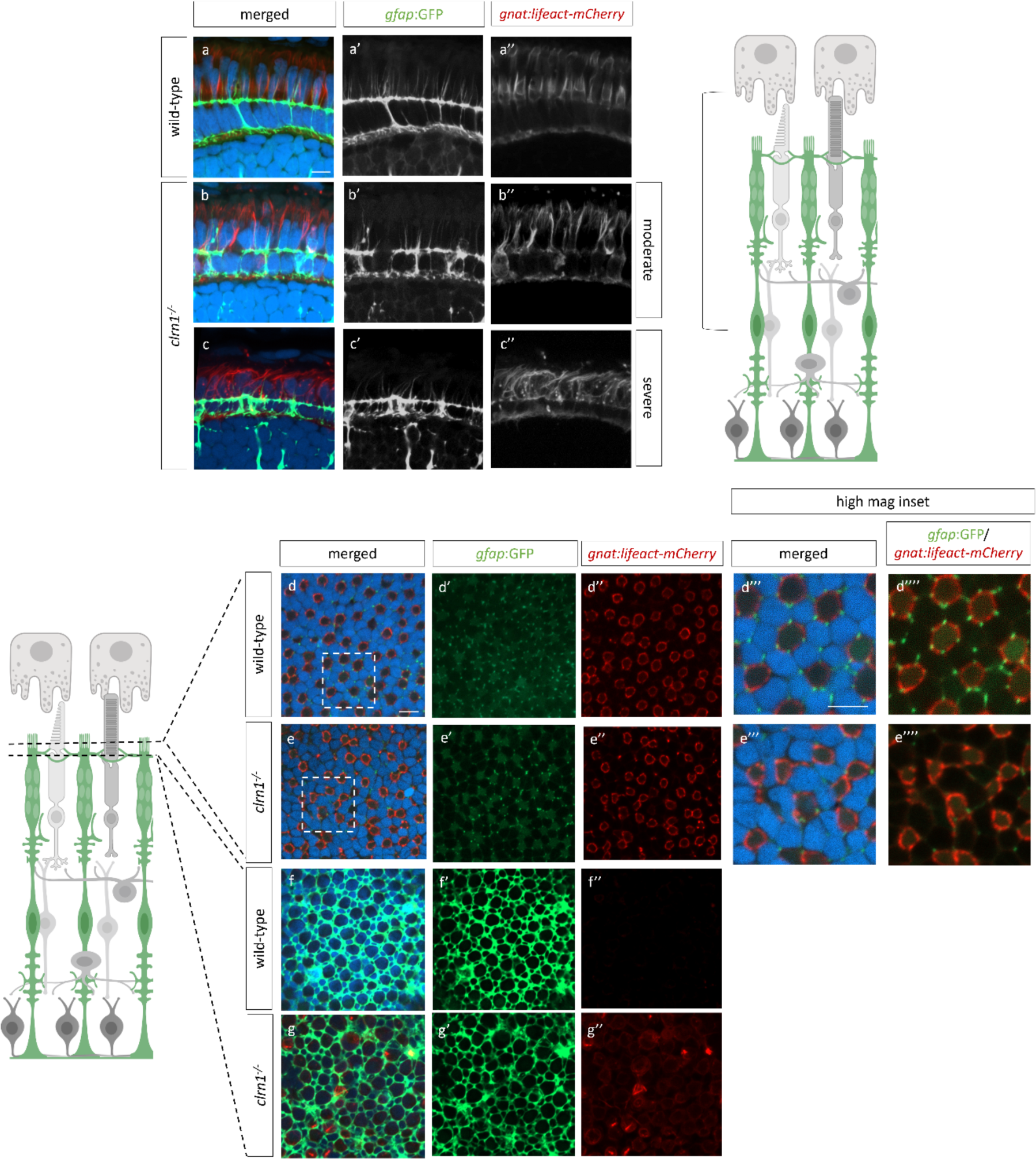
In the absence of Clrn1, cone actin networks and their association with Müller glia apical microvilli are altered. Transverse sections from 7 dpf wild-type (a) and *clrn1^-/-^* Tg(*gfap:gfp*); Tg(*gnat2:lifeact_mCherry*) zebrafish (b,c) highlighting altered Müller glia apical microvilli projections (a’,b’) and alterations in cone-specific actin structures (a’’, b’’). *En face* scan of UV cone outer segment actin structures from wild-type (d) and *clrn1^-/-^*(e) zebrafish at 7 dpf highlighting associations of Müller glia apical microvilli projections (d’, e’) with UV cone actin ring (d’’,e’’). *En face* scans from wild-type (f) and *clrn1^-/-^* (g) zebrafish at the OLM showing Müller glia ring morphology (f’,g’) and differences in the presence of cone actin between wild-type (f’’) and *clrn1^-/-^* (g’’). Scale bar: 5 µm.

*En face* analysis of *Tg(gfap:gfp)* and *Tg(gnat2:lifeact_mCherry)* further highlighted the association of the Müller glia apical projections with calyceal processes of cones, which present as mCherry-positive rings with embedded puncta, highlighting cortical actin and the bundled actin of the calyceal actin. Figure 7d-d’’ shows this structure for the UV cone photoreceptors. In close proximity to each actin ring, we found GFP puncta, which corresponds to the Müller glia apical microvilli. Within each actin ring, there are on average 5-6 associated Müller glia projections. In *clrn1*^-/-^ zebrafish, we observed regions of disrupted actin ring formation (Figure 7e’’). The Müller glia apical projections are also more irregular in their shape and pattern and have reduced association with the cone actin structures.

En face analysis in *clrn1*^-/-^ zebrafish also confirmed altered cone photoreceptor actin and Müller glia association at the level of the OLM. Specifically, we observed regions of actin accumulation, which was not evident in wild-type animals. Analysis of the Müller glia morphology also showed changes in the projections that envelop the photoreceptors at the OLM (Fig 7g, f). These data suggest alteration to the adherens-type junctions of the OLM. Because of these changes, we immunostained for N-cadherin, a component of the OLM junctions. In a previous in vitro study using overexpressed HA-tagged CLRN1, pull-down assays with an anti-HA antibody followed by proteomic analysis identified N-cadherin as one potential CLRN1 binding partner (27). In wild-type animals, as anticipated, we observed an even distribution of N-cadherin staining along the junctions of the OLM (Figure 8a and a’). In *clrn1^-/-^* zebrafish, N-cadherin staining appeared less evenly distributed along the junctions (Figure 8b’). Additionally, we observed a decrease in the localization of N-cadherin at the OLM (Figure 8b’ and d). Outside of the OLM, N-cadherin staining was occasionally mis-localized in a radial pattern extending basally within the mutant retina. (Figure 8b’). Ultra-structural analysis of the OLM junctions revealed that the loss of N-cadherin staining was not due to the failure of junction formation in the absence of Clrn1 (Supplemental Figure 10). Analysis of actin staining, Müller glia morphology, and N-cadherin staining at the OLM does suggest differences in the structural integrity of the OLM. Notably, the OLM highlighted by N-cadherin staining appeared expanded and disorganized. This phenotype is likely due to *clrn1^-/-^* mutants being sensitized to the methanol treatment performed with N-cadherin staining (Figure 8b’), as histology and other marker analyses which do not require methanol treatment did not show this degree of OLM expansion. Quantitation of N-cadherin ring diameter did not highlight any statistically significant differences (p=0.15), but we did observe a shift in the diameter of the N-cadherin rings at the OLM towards a larger size (Figure 8b and e).

**Figure 8.**
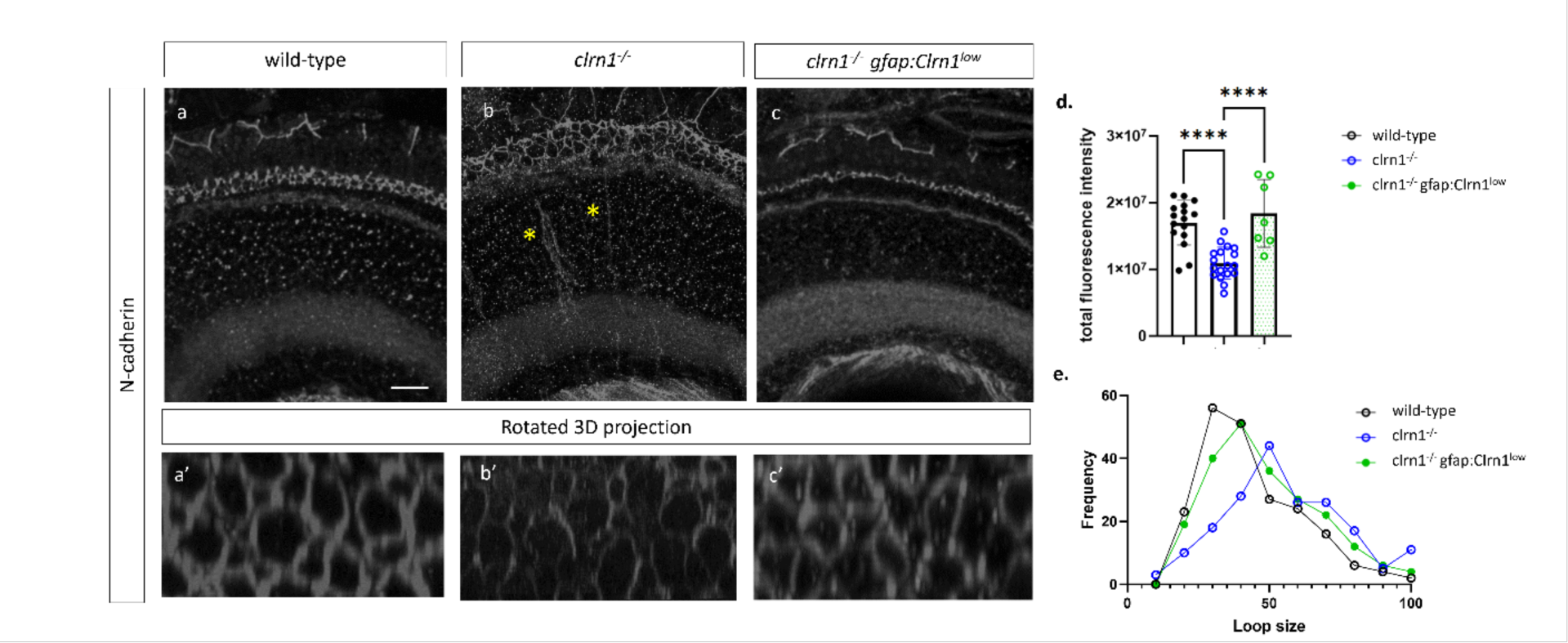
Expression of N-cadherin is reduced in *clrn1^-/-^* zebrafish. (A-C) Transverse cryosections of retinas from 7 dpf wild-type(a) *clrn1^-/-^* (b) and *clrn1^-/-^ Tg(gfap:Clrn1^low^)*(c). Images of N-cadherin staining in the OLM (a’, b’, c’) were generated by rotating 3D rendered Z-stacks taken from the central retina. Quantification of of N-cadherin in a 300 um^2^ region from the central retina revealed that N-cadherin staining was reduced in *clrn1^-/-^* but not in *clrn1^-/-^* gfap*:Clrn1^low^* (D). Plotting of ring diameter frequency plotted as a percentage for 35 rings in an approximately 300 um^2^ region in the central retina. (A-B n:12; C n:7, E n: 6-12) (****p<0.001; One-way ANOVA). Error bars=SD. (A-C) Scale Bar: 10 uM

### Re-expression of Clrn1 in the Müller glia corrects Müller glia projections and N-cadherin localization

Because of prominent Clrn1 expression in Müller glia, we explored whether re-expression of Clrn1 only within Müller glia could rescue the retinal defects caused by its global loss. Specifically, we developed two *Tg(gfap:clrn1-2a-GFP)* lines. The 2a peptide sequence allows for the co-transcription of *clrn1* and *GFP* (Supplemental Figure 8a), facilitating the identification of transgenic zebrafish and cells expressing Clrn1 without hindering Clrn1 function by the addition of an N- or C-terminus epitope (53). Additionally, because *clrn1* and GFP are co-transcribed, the measure of GFP fluorescence intensity allows for relative quantification of transgene expression and, therefore, an indirect estimate of *clrn1* re-expression levels. During the development of the F1 generation, two lines, designated as high re-expression (*gfap*:*Clrn1*^high^) and low re-expression (*gfap*:*Clrn1*^low^) were identified that differed significantly in their level of GFP expression (Supplemental Figure 8b-d). Differences in the expression levels between the *gfap*:*Clrn1*^low^ and *gfap*:CLRN^high^ were quantified on the F2 generation by measuring the total GFP fluorescence intensity (Supplemental Figure 8c). The number of GFP-positive offspring from the F2 generation was also tracked to confirm germline transmission of a single copy for each line.

Using the Müller glia specific Clrn1 re-expression lines, we assessed Müller glia structure in wild-type and *clrn1^-/-^* zebrafish expressing the gfap:*Clrn1*1^high^ or *gfap*:*Clrn1^low^*. For assessing Müller glia structure on the gfap:*Clrn1*1^low^ transgenic lines, *gfap:Clrn1^low^* zebrafish were raised on a *Tg*(*gfap:GFP)* background because the GFP signal from the *gfap*:*Clrn1^low^*transgene was significantly quenched during the process of fixation to levels beyond detection by confocal microscopy (Supplemental Figure 8d). In this instance, a PCR designed towards the *gfap:Clrn1_2a_GFP* sequence was performed to identify zebrafish harboring the *gfap:CLRN1^low^* transgene.

Analysis of Müller glia structure in the *gfap*:*Clrn1*^high^ transgenic zebrafish, in either wild-type or *clrn1^-/-^* mutants, revealed gross alterations in the structure of Müller glia and photoreceptor cells. Specifically, Müller glial process associated with photoreceptor synapses and the OLM were disorganized (Supplemental Figure 9d). In addition, the apical microvilli were reduced – even beyond that of *clrn1^-/-^*mutants alone. Within photoreceptors, actin staining (phalloidin) within the region of the outer segments appeared enhanced and disorganized, likely reflecting the overall structural disruption of photoreceptor outer segments (Supplemental Figure 7d). In *clrn1^-/-^* zebrafish expressing the *gfap*:*Clrn1*^low^ transgene, the Müller glia synaptic and OLM projections, as well as the apical microvilli, showed wild-type morphology (Supplemental Figure 9c). Similarly, within photoreceptors, defects in phalloidin staining were partially rescued. We also observed an increase in the localization of N-cadherin along the junctions of the OLM and an improvement in the distribution of N-cadherin along these junctions within *clrn1^-/-^ Tg(gfap:Clrn1^low^)* zebrafish and partial correction in the N-cadherin ring structure (Figure 8). This analysis overall indicates the importance of appropriate levels of Clrn1 within Müller glia for the structural integrity of the outer retina.

### Targeted re-expression of *clrn1* in Müller glia ameliorates light damage-induced cell death

To determine whether targeted re-expression of *clrn1* in the Müller glia could serve as a therapeutic option for preventing or slowing retinal degeneration in USH3A, we subjected the *gfap*:*Clrn1* transgenic zebrafish to high-intensity light treatment, which was previously established to induce cell death in *clrn1^-/-^* zebrafish (Figure 5). For this experiment, we treated wild-type and *clrn1^-/-^* zebrafish carrying either a *gfap:GFP* control transgene or one of the Clrn1 re-expression transgenes. As before, larvae were exposed to either standard or high-intensity light from 5 to 7 dpf. Several important observations were made from these studies. First, low Clrn1 re-expression within Müller glia of *clrn1*^-/-^ mutants rescued photoreceptor cell death from light stress, as measured by pyknotic nuclei or TUNEL labeling. Low Clrn1 expression in wild-type did not affect wild-type retina. However, high Clrn1 re-expression within Müller glia was damaging to photoreceptors in either normal or light stressed conditions, and in either wild-type or *clrn1^-/-^*mutants (Figure 9). These results both highlight the potential for gene therapy and emphasize the importance of Clrn1 dosage.

**Figure 9:**
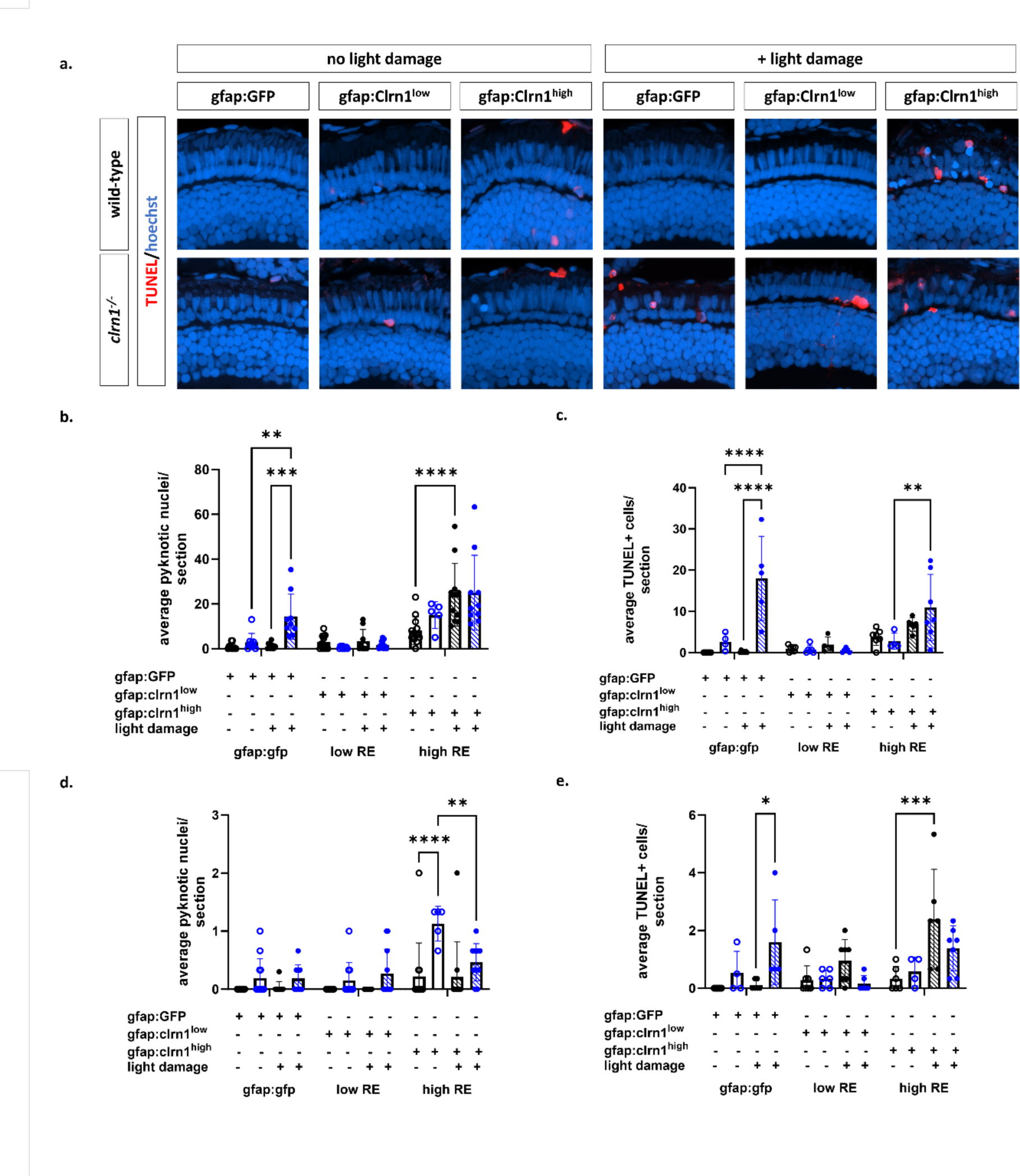
Therapeutic efficacy of Clrn1 re-expression in Müller glia depends on expression level. Animals were treated with 3500 lux for 48 hours from 5-7 dpf (A-L). TUNEL staining on transverse retinal sections from 7 dpf control or light damaged wild-type and *clrn1^-/-^* zebrafish expressing *gfap:GFP*, *Tg(gfap:Clrn1^low^)*, or *Tg(gfap:Clrn1^high^)*. Quantification of pyknotic nuclei in the ONL (M) and INL (O). Quantification of TUNEL positive cells in the ONL (O) and INL (P) reveals that the *Tg(gfap:Clrn1^low^)* transgene decreases sensitivity to light damage induced cell death in *clrn1^-/-^* zebrafish while the *gfap:Clrn1^high^* transgene induced cell death in *clrn1^-/-^* and wild-type zebrafish. Analysis of pyknotic (d) and TUNEL (e) positive cells in the INL of 7 dpf wild-type and *clrn1^-/-^* zebrafish with or without light damage (*p<0.05, **p<0.01, ***p<0.005, ****p<0.001; Two-way ANOVA). Error bars=SD. Scale Bar: 100 µM (n=8-12)

Due to conflicting reports on whether Clrn1 is expressed in the photoreceptor layer (7,25,32–35), we undertook a similar approach to re-express Clrn1 in the rod and cone photoreceptors. We established lines for rod-specific (*Tg*(*rho:clrn1_2a_mCherry*)) and cone-specific (*Tg*(*gnat2:clrn1_2a_mCherry*)) Clrn1 expression. We also established the respective control lines, *Tg*(*rho:mCherry*), *Tg*(*gnat2:GFP*). In contrast to Müller glia re-expression of Clrn1, the rod or cone photoreceptors expressing Clrn1 in a *clrn1^-/-^* mutant background were not protected against light stress (Figure 10).

**Figure 10:**
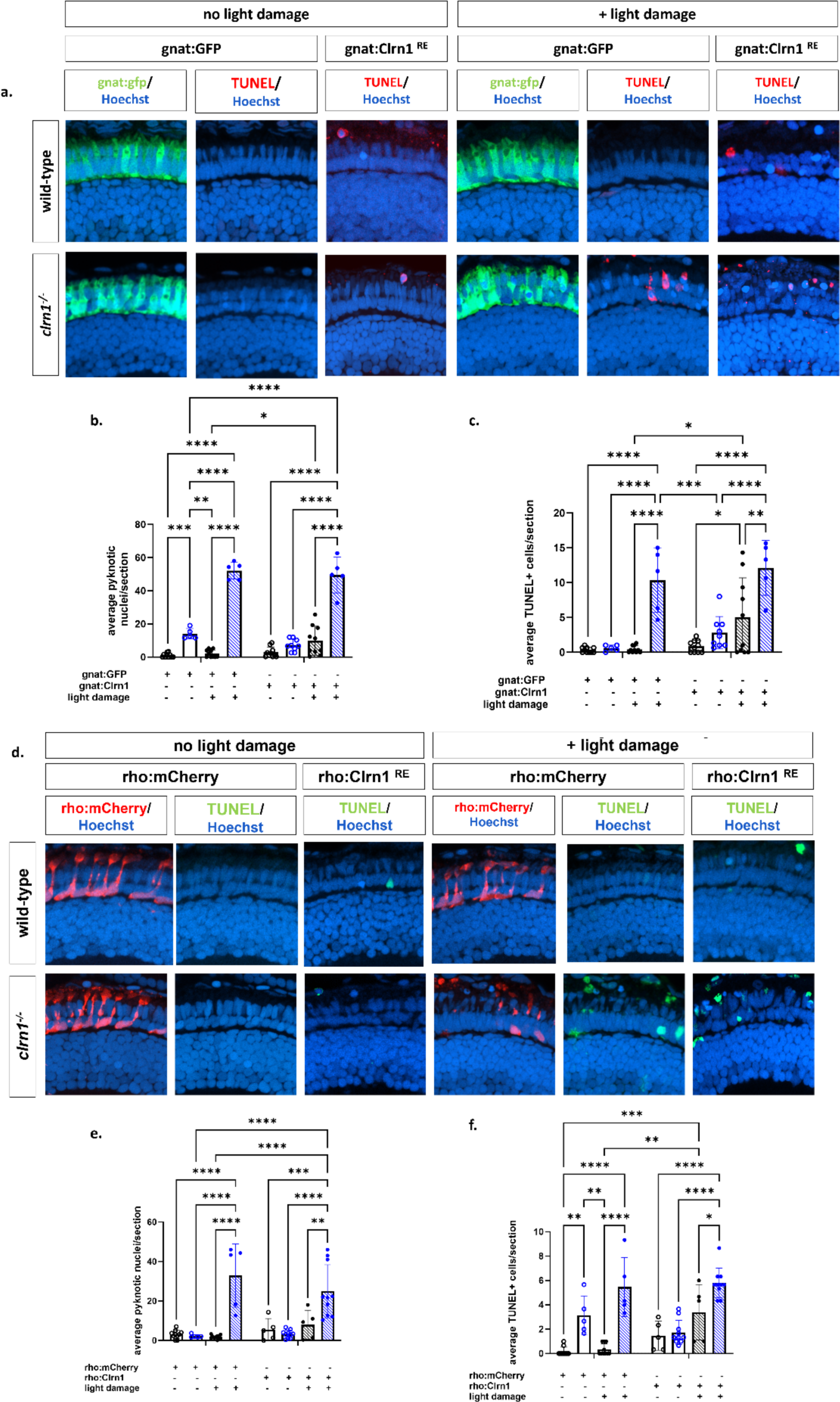
Re-expression of Clrn1 in the rod or cone photoreceptors does not reduce sensitive to light damage. Transverse cryosections from wild-type and *clrn1^-/-^* zebrafish treated with 3500 lux for 48 hours from 5-7 dpf. TUNEL and Hoechst staining on retinal sections from 7 dpf control or light damaged wild-type and *clrn1^-/-^*zebrafish expressing *gnat2:GFP* or *gnat2:Clrn1_2a_mCherry* (a). Quantification of pyknotic nuclei in the ONL (b). Quantification of TUNEL positive cells in the ONL (c). Transverse cryosections from wild-type and *clrn1^-/-^*zebrafish treated with 3500 lux for 48 hours from 5-7 dpf. TUNEL and Hoechst staining on retinal sections from 7 dpf control or light damaged wild-type and *clrn1^-/-^*zebrafish expressing *rho:mCherry* or *rho:Clrn1_2a_mCherry* (d). Quantification of pyknotic nuclei in the ONL(e). Quantification of TUNEL positive cells in the ONL (f). (*p<0.05, **p<0.01, ***p<0.005, ****p<0.001; Two-way ANOVA). Error bars=SD. Scale Bar: 100 µM (n=5-10)

## DISCUSSION

In the current study, we developed a model of USH3A in zebrafish using the CRISPR-Cas9 system to delete the coding sequence of *clrn1*. This represents the first animal model of USH3 that displays retinal degeneration, allowing us to investigate the impact of Clrn1 absence on the retina and the cells that contribute to progressive vision loss. Our results show that USH3A zebrafish retinas are sensitized to high-intensity light, and the cell death observed in the photoreceptor layer was due to loss of both rods and cones. In aging animals, we observed retinal dysfunction at the earliest time point investigated (4 mpf). In larval fish, differential therapeutic potentials were observed following the re-expression of Clrn1 in Müller glia. Low re-expression of Clrn1 in Müller glia was associated with a reduction in photoreceptor apoptosis, underscoring the involvement of Müller glia in the pathophysiology of USH3A. Conversely, with high Clrn1 expression, the therapeutic advantages were negated. Critically, this heightened expression of Clrn1 in Müller glia induced the loss of photoreceptors in wild-type larvae and exacerbated the light stress induced photoreceptor cell death in *clrn1*^-/-^ animals. This data provides important baseline knowledge to guide potential gene therapy initiatives.

Unlike previously described *clrn1* mutant zebrafish, the *clrn1* mutants developed for this study survived into adulthood. Survival differences may be due to the genetic background or husbandry practices, as well as type of mutation. For our allele, we deleted 85% of the coding sequence, resulting in decay of the transcribed mRNA. For the previous USH3A zebrafish model, zinc finger nucleases were used to produce smaller deletions ranging from 7 bp to 43 bp within the first exon of the coding sequence (28). While these mutations lead to the introduction of a premature stop codon, there are potential downstream start sites in subsequent exons, allowing for the potential expression of truncated proteins. Stop codon read-through is also possible. The expression of an altered Clrn1 protein could lead to the activation of stress responses not observed in our *clrn1* knockout zebrafish line. On that note, in humans, one predominant mutation *CLRN1^N48K^* leads to loss of protein glycosylation and improper trafficking of CLRN1 (22,26,27). While we did not observe markers of ER stress in mutant animals (Supplemental Figure 11), the loss of glycosylation and protein trafficking for CLRN1^N48K^ could activate the UPR response, as was previously reported for USH1 proteins in hair cells (54). In support of this idea, genotype-to-phenotype analysis in USH1B patients reports that disease progression is less severe in populations carrying null MYO7A alleles (55).

Consistent with previous reports on *clrn1* mutant zebrafish lines, in the absence of additional stressors, we did not observe any gross histological changes to the structure and lamination of the retina at 7dpf. As others have done, we used high-intensity light treatment to enhance cellular stress on retinal cell types (15–17,56,57). Following light stress, cell death was significantly increased in *clrn1^-/-^* zebrafish, as detected by pyknotic nuclei and TUNEL staining, which were elevated in both rods and cones. Within the peripheral and central retina, we observed a significant reduction in the number of rods and cones in *clrn1^-/-^* zebrafish. While this is the first report of *clrn1^-/-^* zebrafish presenting with a sensitivity to light damage, these findings are in accordance with other zebrafish models of USH (15–17), as well as other retinitis pigmentosa zebrafish models that also presented with increased cell death following treatment with a high-intensity light (58,59).

In non-stressed conditions, we observed progressive functional and structural changes in the retina of *clrn1^-/-^* zebrafish, compared to their age-matched wild-type siblings. At 4 mpf, ERGs revealed rod and cone dysfunction. Interestingly, we did not observe a statistical difference in the average scotopic b-wave, but we did find greater variability in the scotopic B-wave of *clrn1^-/-^* zebrafish. Specifically, 30% of the animals presented with a hyper-response. At this same age, we observed an increase in BrdU-positive cells, which predominantly occurred in the region of the retina where the rod nuclei reside. Newly regenerated photoreceptors may affect the B-wave due to their immaturity or may promote circuit re-wiring as they integrate into the existing retina. At 4 mpf, we also observed a significant decrease in the photopic B-wave, implicating cone dysfunction. The decrease in the photopic ERG response could be due to the degeneration of cones or impaired cone function. Alternatively, Müller glia defects may affect opsin recycling, and account for differences in the scotopic and photopic responses. In mice, scotopic b-wave enhancements were observed during light-adapted ERGs early in the degenerative process of Rd10 (PDE6 mutant) mice (60). In patients with cone dystrophy with supernormal rod response, patients present with elevated rod b-wave responses at higher light intensities, which is in agreement with the observations of hyper-responding *clrn1^-/-^*zebrafish (61,62). However, the ERG presentation in *clrn1^-/-^* zebrafish deviates from the ERG recordings described for USH3A patients. In patients with USH3A, early ERG dysfunction begins with a reduced scotopic response and progresses to reduced photopic response, as is typical of rod-cone dysfunction. The species differences in progression of ERG defects may be due to several reasons. These include (A) the more heterogenous rod-to-cone composition across the human retina compared to the consistent ratios in zebrafish, (B) potential activation of regeneration in zebrafish, or (C) differential roles of Müller glia in retinoid recycling.

Towards identifying a potential retinal function of *clrn1* and the pathological factors promoting retinal degeneration in USH3A, we focused on characterizing alterations in the actin-based structures in the outer retina. We found that the loss of Clrn1 leads to changes in the Müller glia apical microvilli and contacts between photoreceptors. The apical microvilli projections appear more disorganized, with variable lengths, similar to reported stereocilia phenotypes of the inner ear of *clrn1* mutants (63). These results are consistent with the function of Clrn1 in actin organization and dynamics. In wild-type fish, the Müller glia projections appeared to envelop the base of the photoreceptor outer segments, suggesting structural support. Similar findings have also been recently highlighted by Sharkova *et al* (10). We further described a close association between the Müller glia apical microvilli and cone calyceal processes, a discovery facilitated through our development and application of the cone-specific Lifeact transgenic line (Figure 7). Our analysis of *clrn1*^-/-^ mutants suggests that the association between Müller glia apical microvilli and photoreceptor calyceal processes is mediated by Clrn1 (Figure 11). We anticipate this may be through direct or indirect interaction between Muller glia expressed Clrn1 and photoreceptor expressed USH-associated proteins. In accordance with this hypothesis, the enhanced cell death we observed following high-intensity light stress may result from the reduced structural integrity. Additionally, alterations at the *clrn1* mutant OLM, including N-cadherin distribution are consistent with another role of Müller glia-derived Clrn1 in mediating the integrity of the adherents-type junctions and thus providing additional structural support to photoreceptor (Figure 11). Clrn1 function at the OLM may in part be mediated through interactions with N-cadherin, and Müller glia expressed USH-associated proteins, such as Harmonin. In accordance with this, defects observed in our *gfap*:Clrn1high transgenic animals; may in part be a consequence of perturbing secondary functions of Clrn1 interactors; such as Harmonin an essential protein in photoreceptor synaptic development (14). Accordingly, further work identifying Clrn1 interactors will be essential in conferring changes directly induced by the loss of Clrn1 function versus those occurring as a secondary consequence.

**Figure 11:**
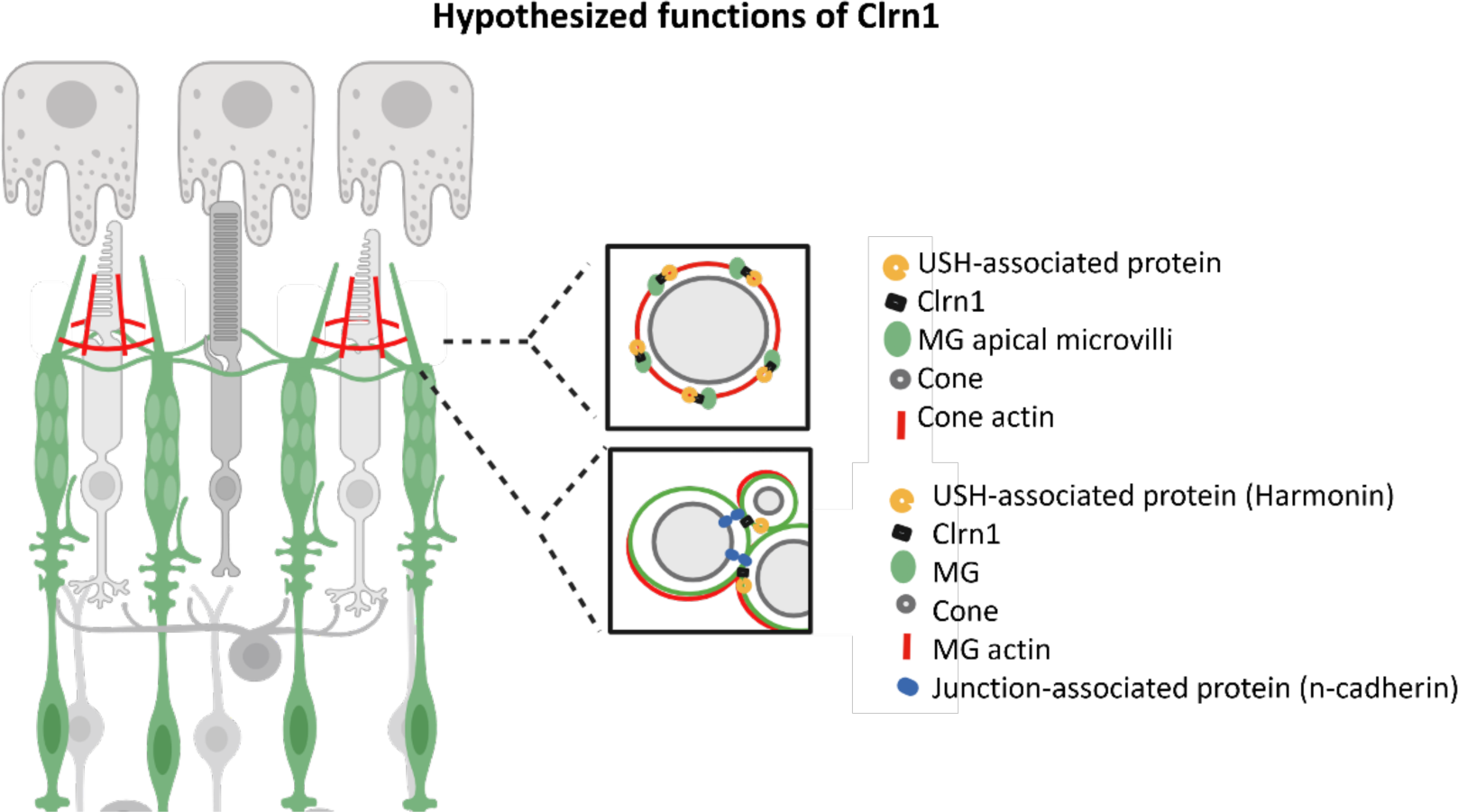
Hypothesized function of Müller glia expressed Clrn1. Based on our findings, we propose that Clrn1 is important for actin-based Muller glia structure(s) in the outer retina. Our results suggest a model in which Clrn1 localization within Muller glia apical projections facilitates the interaction between these structures and photoreceptor calyceal process. In addition, results suggest that Clrn1 also has a role at the OLM to regulate appropriate composition of adherens junction proteins. These proposed functions may be necessary to provide structural support for photoreceptors to appropriately respond to mechanical stress experienced within the retina.

Overall, our results suggest that Clrn1 within Müller glia has roles in regulating the structure of Müller glia and non-autonomously maintaining photoreceptor homeostasis. Future studies aimed at the identification of Clrn1-binding partners within Müller glia could assist in better defining precise cellular mechanisms. Importantly, these findings suggest gene replacement therapy for USH3A patients holds promise, but the dosage and timing of CLRN1 expression within Müller glia must be carefully evaluated.

## MATERIALS AND METHODS

### Zebrafish Maintenance

All transgenic and mutant lines were generated and maintained in the ZDR genetic background. When possible and if otherwise not noted, wild-type siblings or cousins of each line were used as control groups. Zebrafish (*Danio rerio*) were maintained at 28.5°C on an Aquatic Habitats recirculating filtered water system (Aquatic Habitats, Apopka, FL) in reverse-osmosis purified water supplemented with Instant Ocean salts (60mg/l) on a 14 h light: 10 h dark lighting cycle and fed a standard diet. All animal experiments were approved by the Institutional Animal Care and Use Committee of the Medical College of Wisconsin.

### Generation of *clrn1* mutant zebrafish

Due to the small size of *clrn1* (ENSDART00000171202.2), 85% of the coding sequence was deleted using a gRNA targeting Exon1 (5’-CTCGACTCAGTTTTGGGTTCAAGC-3’) and another gRNA targeting the 3’UTR (5’-AGCTCGCTTAAGCCATTGAGAGCA-3’). gRNAs were purchased from IDT (Coralville, IA) and processed according to manufacturer instructions. For the generation of *clrn1* mutant lines, gRNAs (10 ng/µl) and Cas9 protein was co-injected into 1-to 4-cell zebrafish embryos from wild-type ZDR fish maintained internally in the Link lab., respectively. Surviving embryos were raised to adulthood before outcrossing to identify the founder fish carrying germline edits in *clrn1.* Offspring from these fish were raised to adulthood, thereafter, fin-clipped for genotyping. The resulting deletion was confirmed via sequencing (Retrogen, San Diego, California, USA).

### Genotyping

Genomic DNA was extracted from zebrafish tissue using a Puregene Core Kit (Qiagen, Germantown, MD). The genomic region containing the desired large deletion was amplified using primers external to each cut site. For detection of the mutant band the forward primer sequence used 5’-GTTCATATATTCAGGCGTGC-3’ and the reverse primer sequence used was 5’-AGAGGAAACTGTGATGTCCC-3’. The thermocycler conditions for detecting the presence of a large deletion allele were designed using extension times, allowing only the amplification of the mutant region. When detecting the presence of a wild-type allele a third internal primer located within the sequence deleted was. The sequence for the internal reverse primer was 5’-AGTTGGGTTTAGGTTTGGGTAG-3’. For genotyping *Tg(gfap:Clrn1^low^)* larvae on a *Tg(gfap:GFP)* background, the following primers were used: 5’CGCCTGCTTGGTCCTCATTT-3’ and 5’-ACAGAGAGAAGTTCGTGGCT-3’.

### RNA extraction and amplification of *clrn1* cDNA

For mRNA isolation, samples were homogenized in Trizol (ThermoFisher, Waltham, MA) followed by mRNA extraction using Trizol-chloroform treatment. The isolated aqueous phase from the resulting Trizol-chloroform phase separation is transferred to a new tube and incubated with isopropanol and glycogen at room temperature for 10 minutes. Samples were centrifuged at 4°C for 10 minutes to pellet the precipitated RNA. Isolated RNA was washed with 75% ethanol in DEPC-treated water. Following this wash, the pellet was briefly dried then resuspended in DEPC-treated water with a 10-minute incubation at 60⁰C. Resuspended RNA is then subjected to a DNase I treatment, and concentration is quantified. cDNA was generated using the Superscript III First-Strand Synthesis System for RT-PCR Kit (Invitrogen, Waltham, MA) per manufacturer’s instructions and all qRT-PCR was performed on a CFX Connect Real-Time System (Bio-Rad, Hercules, CA) using prime time gene expression master mix (IDT).

### Plasmid Construction

Plasmids were constructed using Gateway assembly (Thermo Fisher Scientific, Waltham, MA). The pME’ *clrn1* entry clone was generated by amplifying the full-length cDNA of *clrn1* (ENSDART00000171202.2) with primers containing attB recombination sites (Clrn1 attB2 FPrimer: 5’-GGGGACAAGTTTGTACAAAAAAGCAGGCTCCATGCCTAACCGTCAAAAGCA-3’ Clrn1 attB3 FPrimer: 5’-GGGGACCACTTTGTACAAGAAAGCTGGGTCGTACATGAGATCTGCAGCTC -3’) and placed into a pME’ entry clone. The *gfap:clrn1-2a-GFP*, *rho:clrn1-2a-mCherry*, and *gnat2:clrn1-2a-mCherry* plasmids were generated using the three-part Gateway system. Specifically, the *pME’ clrn1* was recombined with respective cell-specific p5E’ plasmids (Müller glia: *gfap*, rod photoreceptors: *rho*, cone photoreceptors: *gnat2*) and p3E’ 2a-GFP. The backbone vector contained Tol-2-inverted repeats flaking the transgene construct, which were used to facilitate plasmid insertion into the zebrafish genome (Kwan et al., 2007). Similarly, the control plasmids (*gfap:GFP*, *rho:mCherry*, and *gnat2:GFP*) were generated using a p5E’gfap promoter, pME’ eGFP or pME’mCherry, and p3E’ polyA into a backbone vector with Tol-2 inverted repeats. For analysis of cone photoreceptor actin structures, a pME-Lifeact (65) plasmid was a gift from Rob Parton (Addgene plasmid #109545). Using gateway cloning, this was recombined with a p5E’ *gnat2* promoter and p3’ mCherry into a backbone vector with Tol-2 inverted repeats.

### Generation of transgenic lines

To generate transgenic zebrafish, transposase mRNA (10 ng/ul) was injected with plasmid DNA (10 ng/µl) to generate F0 transgenic lines. F0 fish were screened for reporter fluorescence in the Müller glia. Expressing F0 fish were raised to adulthood and outcrossed to establish F1 transgenic zebrafish with germline integration of the transgene. F1 transgenic larvae were raised to adulthood and outcrossed to wild-type zebrafish. Clutches with around 50% of F2 embryos with reporter expression were identified, suggesting a single active copy of the transgene were used for subsequent analysis.

### Immunofluorescence

To prepare larvae for cryosections, animals were anesthetized with 0.016% tricaine methanesulfonate and fixed overnight at 4°C in 4% PFA. Following fixation, larvae were washed three times for 10 minutes with PBS. Larvae were subsequently stepped through a sucrose gradient and incubated in OCT compound (Optimum Cutting Temperature medium) overnight. Following overnight incubation in OCT compound animals were mounted in OCT compound, frozen on dry ice, and then stored in the -80°C until sectioned. For staining, retinal cryosections were treated with PBST (PBS with 0.1% Triton 100) for one hour, then incubated in blocking solution containing 5% normal goat serum for one hour. Primary antibody was diluted in PBST containing 5% normal goat serum (NGS), and sections were incubated overnight at 4°C. Following washes in PBST, sections were stained with secondary, diluted in PBST containing 2% normal goat serum. For phalloidin staining, sections or whole mount preparations were pre-incubated in PBST for one hour. Samples were stained for one hour with phalloidin Alexa fluor 488 (1:300, ThermoFisher, A12379) or phalloidin Alexa fluor 568 (1:300, ThermoFisher, A12380). For PNA and WGA staining, animals were stained before sectioning. Specifically, animals were treated with PBST for one hour, followed by 4 hours of staining with rhodamine conjugated PNA (1:250, Vector Labs RL 1072) and WGA alexa fluor 488 (1:250, ThermoFisher, W11261) or WGA alexa fluor 594 (1:250, ThermoFisher, W11262). Larvae were then washed with PBS and processed as discussed above. For all staining, samples were counter stained with Hoechst (1:1000, ThermoFisher, 62249). Slides were cover slipped with Vectashield (Vecto Labsr, H-1000) and analyzed by confocal microscopy.

### Paraffin Histology

Adult fish used for paraffin histology were fixed in 10% neutral buffer formalin overnight at 4°C. Prior to fixation, heads were bisected to allow for better fixation of the retina. Samples were then mounted in histogel and processed in paraffin on a Sakura VIP5 automated tissue processor (Sakura Finetek Europe, Flemingweg, The Netherlands) for histology and immunohistochemistry. After paraffin embedding, samples were sectioned at 4 μm (Microm HM355S, ThermoFisher Scientific, Waltham, Massachusetts, USA) onto poly-l-lysine coated slides and air-dried at 45°C overnight for any subsequent immunohistochemistry or routine H&E staining.

### Immunohistochemistry - paraffin sections

All slides were dewaxed prior to their optimal antigen retrieval protocol. All antibodies used a citrate buffer epitope retrieval (DAKO, Agilent). Slides were washed in PBST to remove excess retrieval buffer. Slides were blocked for one hour in PBST with 5% NGS. Samples were incubated with primary antibodies overnight at 4°C. After primary, slides were washed in PBS, then incubated with secondary antibodies (1:500) diluted in PBST with 2% NGS. Slides were washed in PBS then counterstained with Hoechst (1:1000, ThermoFisher, 62249) for 15 min. Sections were protected with Vectashield mounting medium (Vector Labs, H-1000).

### RNAscope *in situ* hybridization

*In situ* hybridization (ISH) was performed on paraffin-embedded zebrafish retina sections. *Clrn1* transcripts were detected using an automated Leica Bond platform with heat-induced Leica ER2 antigen retrieval buffer solution and RNAscope 2.5 LS Protease III digestion (Advanced Cell Diagnostics, Hayward, CA, USA), as previously described (32). Briefly, after hybridization to 20ZZ probes targeting the region comprising nucleotides 2-911 of Dr-clrn1 NM_001002671.1, a six-step amplification process was performed, followed by chromogenic detection using Fast Red (Advanced Cell Diagnostics). Images were collected with a fully automated widefield DMi8 Leica fluorescence microscope.

### Antibodies

Anti-BrdU was purchased from Abcam (Rat monoclonal, ab6326) and used at a dilution of 1:300 for the staining of paraffin sections. The anti-SV2 antibody was purchased from Developmental Studies Hybridoma Bank (mouse monoclonal, SV2s) and used at a dilution of 1:100 for the staining of paraffin sections. Anti-CTBP2 (Ribeye) was purchased from Proteintech (Rabbit polyclonal, 10346-1-AP). The secondary antibodies use for immunofluorescent detection were goat anti-rat secondary Alexa fluor 488 (ThermoFisher A-11006), goat anti-rabbit secondary Alexa fluor 568 (ThermoFisher, A-11011), goat anti-mouse secondary Alexa fluor 568 (ThermoFisher A-32742).

### Fluorescent Microscopy

Confocal microscopy was performed using a Nikon Eclipse E800, C2 Nikon Eclipse 80i, or Zeiss LSM 980 confocal microscope. For whole mount imaging, larvae were embedded in 1% low-melting agarose in glass-bottomed Petri dishes. Images were generated using ImageJ software (Rasband, W.S. ImageJ, U.S. National Institutes of Health, MD).

### Spectral Domain – Optical Coherence Tomography (SD-OCT)

Zebrafish eyes were imaged using a Bioptigen Envisu R2310 SD-OCT imaging system, equipped with a 12 mm telecentric lens (Bioptigen, Morrisville, NC) using a Superlum Broadlighter T870 light source centered at 878.4 nm with a 186.3 nm band width (Superlum, Cork, Ireland). Volume scans were nominally 1.0 × 1.0 mm with isotropic sampling (500 A scans/B scan; 500 B scans). Raw OCT scans for retinal images were exported and processed using a custom OCT volume viewer (Java software, Oracle Corporation, Redwood Shores, CA), in which an adjustable contour is used to generate *en face* summed volume projection (SVP) images (66,67). For a given B-scan, 15 control points were added to the initial contour, where each control point is adjusted to follow the contour of the layer(s) of interest. Multiple *en face* images can be generated for each OCT volume, resulting in images of different retinal features (*e.g*., inner retinal vasculature and photoreceptor mosaic). *En face* images of the NFL, RPE, and UV cones were generated to visualize the cone mosaic, RPE pigmentation, and gaps in the reflectivity of the NFL. Using *en face* images of the UV cone mosaic, mosaic geometry were assessed from the resultant cone coordinates using a custom program as previously described (39).

### OMR

The OMR assay was performed on 7dpf larvae that were maintained in standard lighting conditions or treated with high-intensity light from 5-7 dpf. To test for OMR, larvae were placed in a 48-well plate placed on top of a tablet playing the stimulation videos. Animals were recorded for a three-minute period with direction changes at 1-minute intervals. Animals were scored on their ability to detect direction changes within the first 10 seconds. Zebrafish that could not be tracked for all three passes were not included in the analysis. Stimulation used for assay was generated using publicly available software developed by Brastrom *et al* (68).

### BrdU Treatment

To assess for regeneration, BrdU labeling was performed to label newly generated cells in the adult zebrafish retina. Prior to collection, adult animals were placed in fish water containing 10 mM 5-bromo-2’-deoxyuridine (BrdU) (Abcam, Cambridge, UK) overnight 1 and 2 weeks before animals were sacrificed. After treatment, animals were rinsed twice for 10 min in fresh fish water to rinse off excess BrdU.

### TUNEL Staining

TUNEL technique using the *in-situ* cell death detection kit, Roche (Millipore Sigma CA, USA), was performed according to the manufacturer’s instructions on cryosectioned samples. For staining, slides were incubated in PBST for one hour (PBS with 1% TritonX-100). Following PBST, the samples were incubated at 37°C with the TUNEL reaction mixture (containing 5 μl of TdT + 45 μl of fluorescein conjugated dUTP) for two hours. Following incubation in TUNEL mixture, samples were washed for 30 minutes in 1X PBST. Slides were co-stained with Hoechst for 15 min, then washed 3 times for 10 min in PBST. The number of TUNEL positive cells were manually counted and averaged across three sections in the central retina.

### Histology and Transmission Electron Microscopy

7 dpf larvae were fixed with 1.0% paraformaldehyde, 2.5% glutaraldehyde in 0.06-M cacodylate buffer (pH 7.4) overnight at 4°C. Post fixation, samples were washed in cacodylate buffer and post-fixed with 1% osmium tetroxide and then dehydrated by a series of methanol and acetonitrile washes. Larvae were infused with Epon 812 resin (Electron Microscopy Sciences, Hatfield, PA, USA) incubating with 1:1 acetonitrile:Epon for one hour, followed by an incubation in 100% Epon in a 37°C degree water bath, then 100% Epon in a 37°C heat block. Finally, larvae were embedded in 100% Epon and hardened at 65°C for 24 hours. For transmission electron microscopy (TEM) analysis, 70-nm sections were cut, collected on hexagonal grids, and stained with uranyl acetate and lead citrate for contrast, followed by imaging on a Hitachi H-600 Transmission Electron Microscope (Hitachi, Ltd., Tokyo, Japan).

### *Ex Vivo* Electroretinograms

*Ex vivo* ERGs were performed using the OcuScience (Henderson, Nevada) *Ex* Vivo ERG adapter according to previous published protocols, with appropriate modifications for analysis of zebrafish retinal function (69). Prior to performing ERGs, animals were dark adapted overnight, and all experimental setups were performed under dim red illumination. Zebrafish were anesthetized in 0.016% tricaine methanesulfonate. Once anesthetized, optic cups were dissected in AMES media (A1420, Sigma-Aldrich, CA, USA). To dissect optic cups, eyes were punctured with a 28-gauge needle. Two forceps were used to peel off the sclera and RPE. During this process the lens was also removed. Samples were then mounted in the *ex vivo* ERG adaptor.

During recording, samples were perfused with AMES media at a rate of 5-10 ml/hr. ERG recordings were carried out using an Espion E2 system (Diagnosys LLC, Cambridge, UK) with a low frequency filter of 0 Hz, high frequency filter of 1000 Hz, and notch filter of 60 Hz (Bessel filters). The scotopic session included a single flash stimulus increasing from 0.1 mcds/m^2^ to 25 cds/m^2^. Six response per intensity level were averaged, with an inter-stimulus interval of 5 seconds for stimuli ranging from 0.1 mcds/m^2^ to 100 mcds/m^2^. For stimuli above 100 mcds/m^2^, an inter-stimulus interval of 17 seconds was used. For the photopic session animals were light adapted for 10 min with a background illumination of 30 cd/m^2^. Following the light adaptation of 10 min, the photopic session began with single flash recordings as above but with a reduced number of light intensities (0.03, 0.1, 1, 3, 10 and 25 cds/m^2^). Subsequently, flicker ERGs were obtained with flashes of 3 cds/m^2^ using frequencies of 1, 2, 5, 10, 15, 20, 25 and 30 Hz. During recording, all equipment was located within a Faraday cage to minimize external electrical noise.

### Statistical analysis

Mean or total pixel intensity was measured using ImageJ (Rasband, W.S. ImageJ, U.S. National Institutes of Health, MD). Data was processed using Microsoft Excel (Microsoft, Redmond, WA) and graphed using GraphPad Prism (GraphPad, La Jolla, CA). An unpaired, two-tailed t test was used to analyze graphs with two groups. For three or more groups, a one-way or two-way ANOVA was conducted with Tukey’s post-hoc analysis for pair-wise comparisons.

## ACKNOWLEDGMENTS

Supported by funding from National Institutes of Health grant R01 EY026559-06A1 (AD, BAL), 5R21EY032575 (AD), Foundation Fighting Blindness awards TA-NMT-0621-0811-UFL (AD) and PPA-0617-0718 (BAL), and The E. Matilda Ziegler Foundation for the Blind, Inc. (RFC). Last, components of this investigation were conducted in a facility constructed with support from Research Facilities Improvement Program, Grant Number C06RR016511, from the National Center for Research Resources, National Institutes of Health.

## FIGURES AND LEGENDS

**Supplemental Figure 1:**
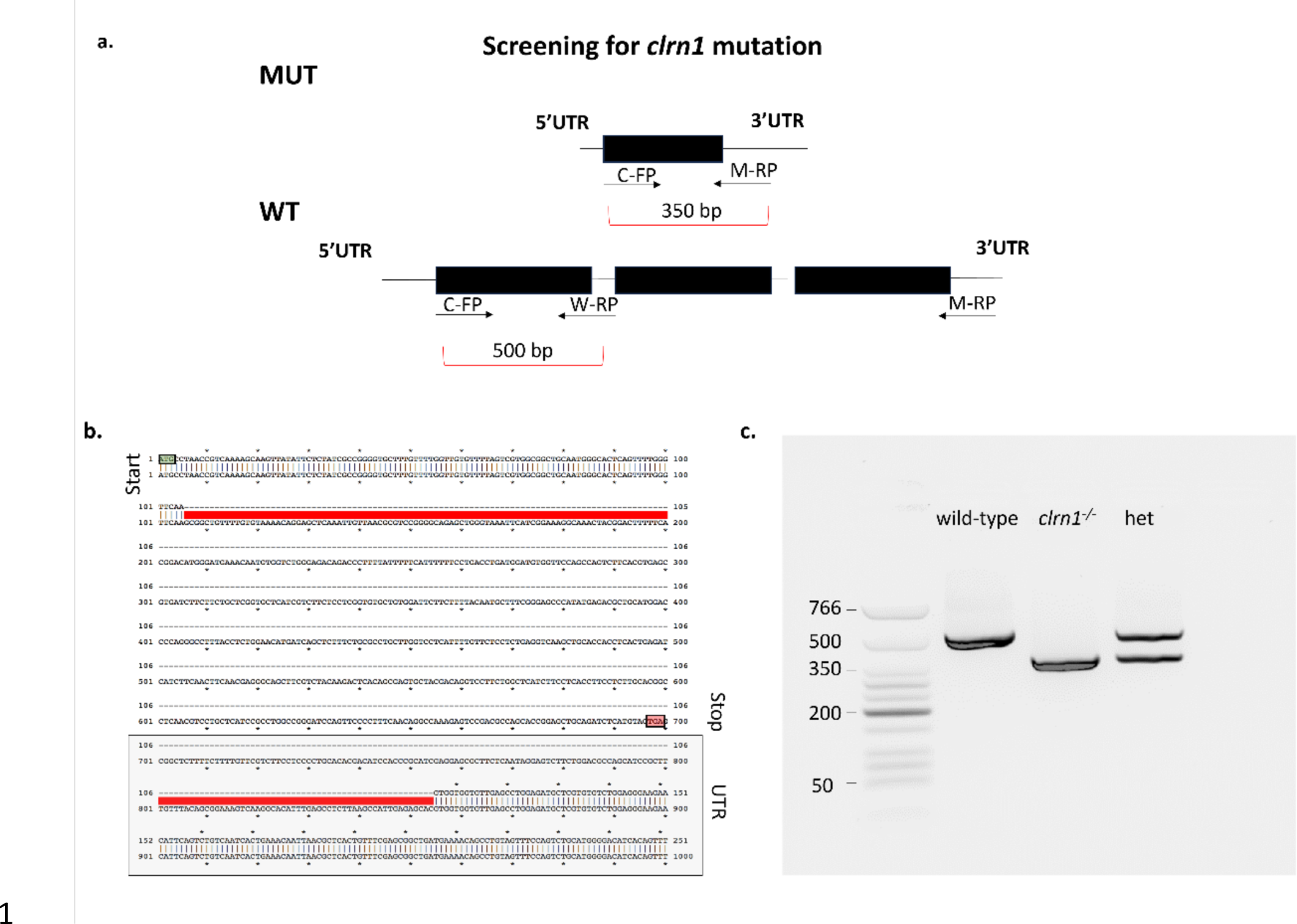
Method to genotype for *clrn1^-/-^*, wild-type, and heterozygous zebrafish. (a) Depiction of primer design for genotyping wild-type, heterozygous, and *clrn1^-/-^* zebrafish. A common forward primer (C-FP) was designed upstream of the exon 1 cut site, while a mutant reverse primer (M-RP) was designed downstream of the 3’ UTR cut site, and W-RP was designed internal to the exon 1 cut site. (b) Generation of the large deletion in the *clrn1* coding sequence was confirmed with Sanger sequencing. (c) Example gel depicting the PCR amplification products for wild-type, *clrn1^-/-^*, and heterozygous (het) zebrafish.

**Supplemental Figure 2:**
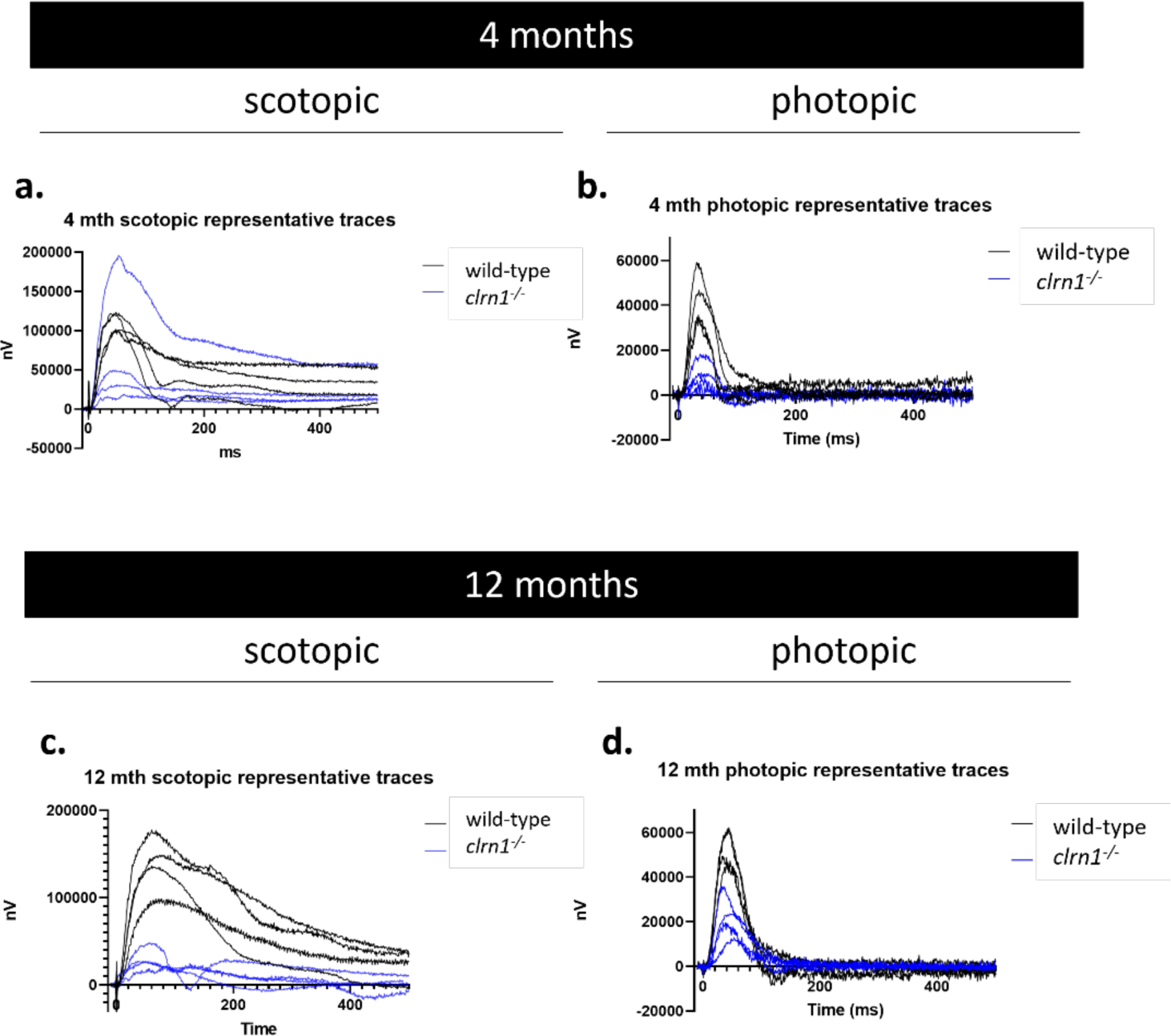
Representative traces of photopic and scotopic responses. Representative traces from four 4 mpf wild-type and *clrn1^-/-^* scotopic (a) and photopic (b) responses to highlight high variability in the *clrn1^-/-^* scotopic b-wave. Representative traces from 12 mpf wild-type and *clrn1^-/-^* scotopic (c) and photopic (d) responses.

**Supplemental Figure 3.**
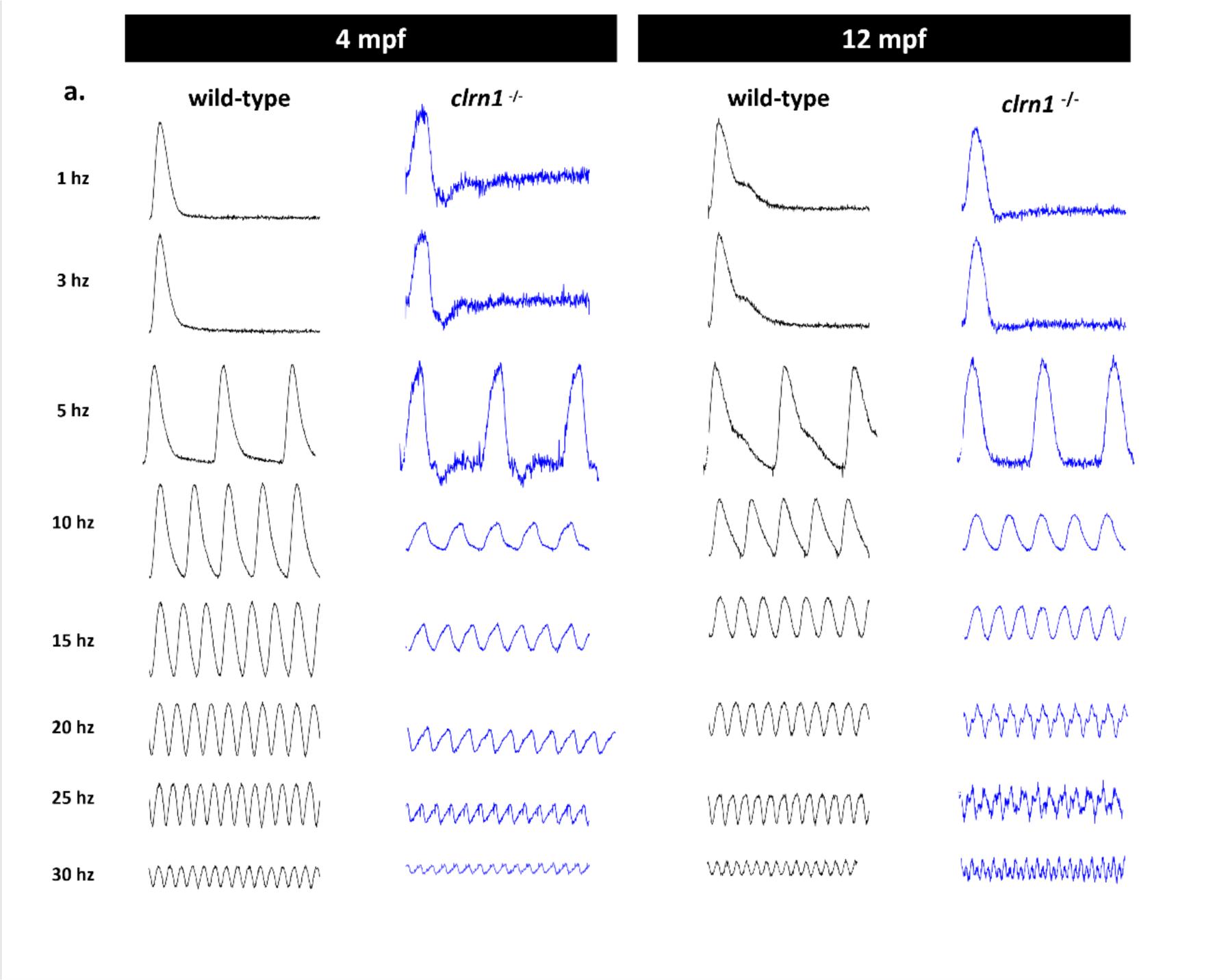
Photopic flicker response are reduced in 4 and 12 mpf *clrn1^-/-^*zebrafish. (a). Representative photopic flicker traces of 4 and 12 mpf wild-type and *clrn1^-/-^* at a range of 1-30 hz.

**Supplemental Figure 4.**
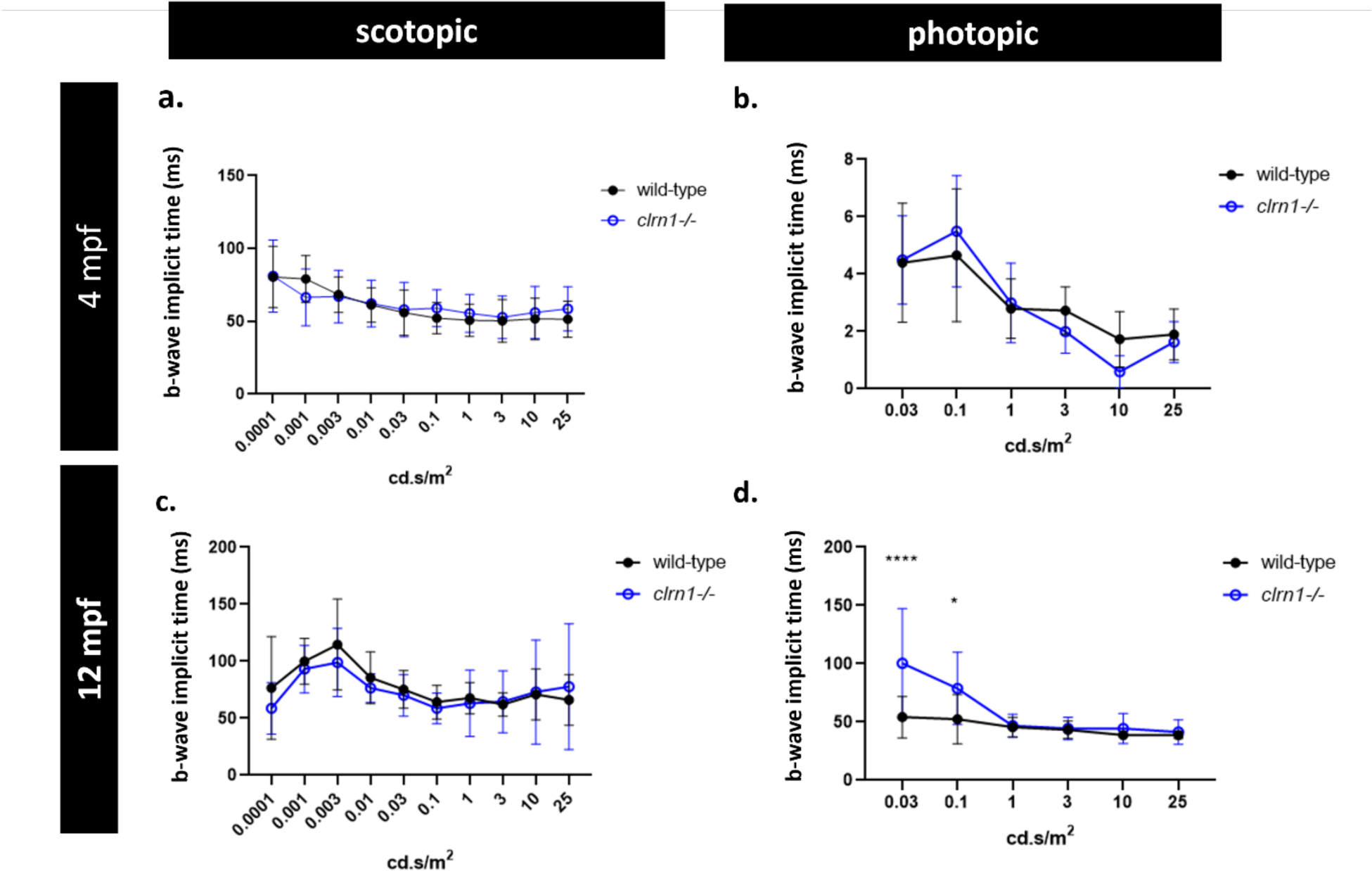
Scotopic and photopic B-wave implicit times at 4 and 12 mpf. Scotopic (a) and photopic (b) b-wave implicit time for wild-type and *clrn1^-/-^*zebrafish at 4 mpf. Scotopic (c) and photopic (d) b-wave implicit time for wild-type and *clrn1^-/-^* zebrafish at 12 mpf. (*p<0.05, ****p<0.001; One-way ANOVA) (n=10 per group). Error bars=SD.

**Supplemental Figure 5:**
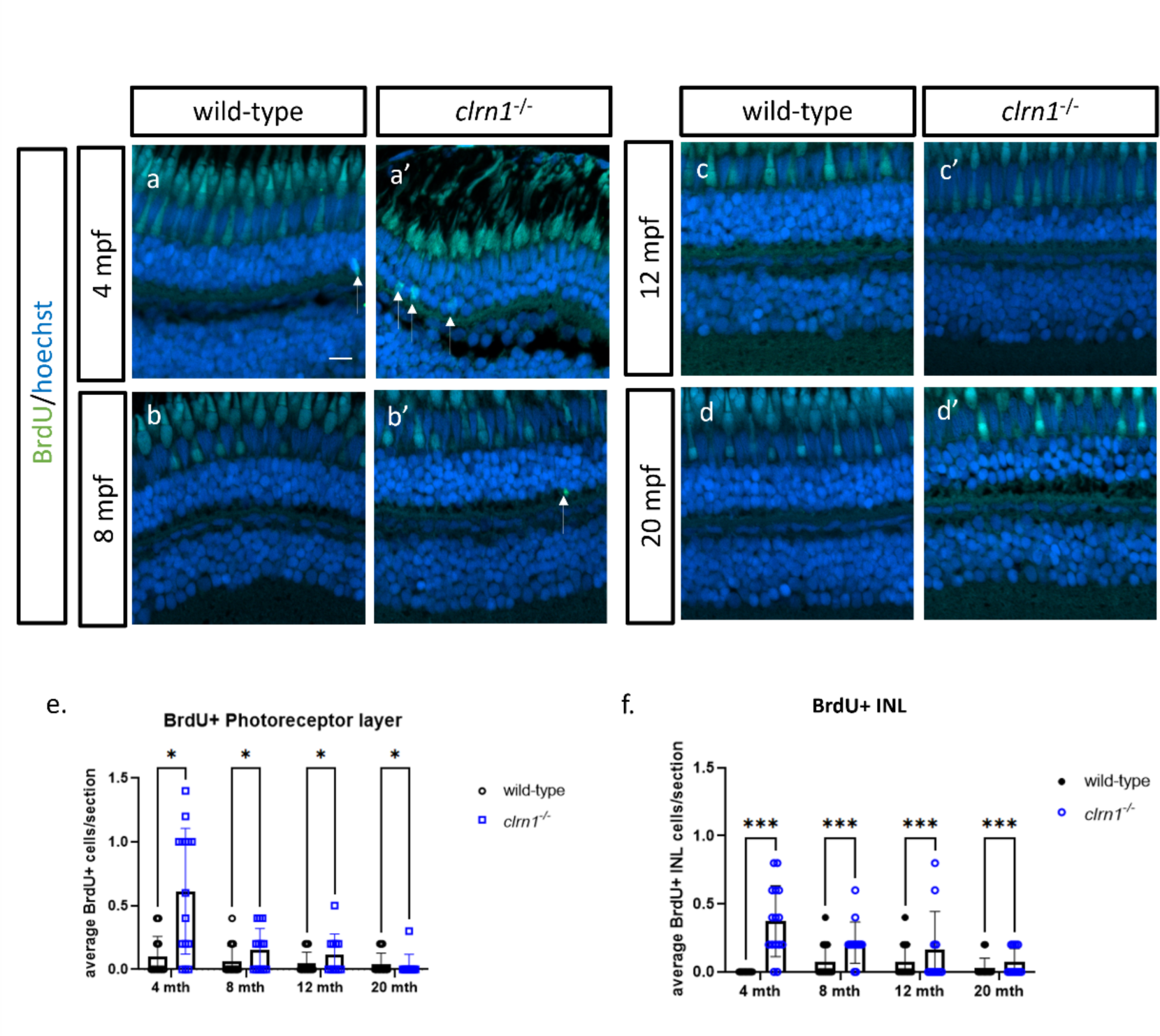
BrdU incorporation is elevated in *clrn1^-/-^* zebrafish at 4 mpf and decreases with age. (a-d) Anti-BrdU (green) staining marks cells with BrdU incorporation, which serves as a measure of regeneration. Paraffin sections from 4 mpf (a), 8 mpf (b), 12 mpf (c), and 20 mpf (d) wild-type and *clrn1^-/-^* zebrafish retinas. Quantification of Brdu+ nuclei in the photoreceptor (PR) (e) and inner nuclear layer (e) revealed an increase in BrdU incorporation for *clrn1^-/-^* zebrafish at the youngest time point and this decreased with age. White arrows highlight BrdU+ nuclei. (**p<0.01; One-way ANOVA). Scale bar: 10 µm. White arrows denote Hoechst and BrdU positive nuclei.

**Supplemental Figure 6:**
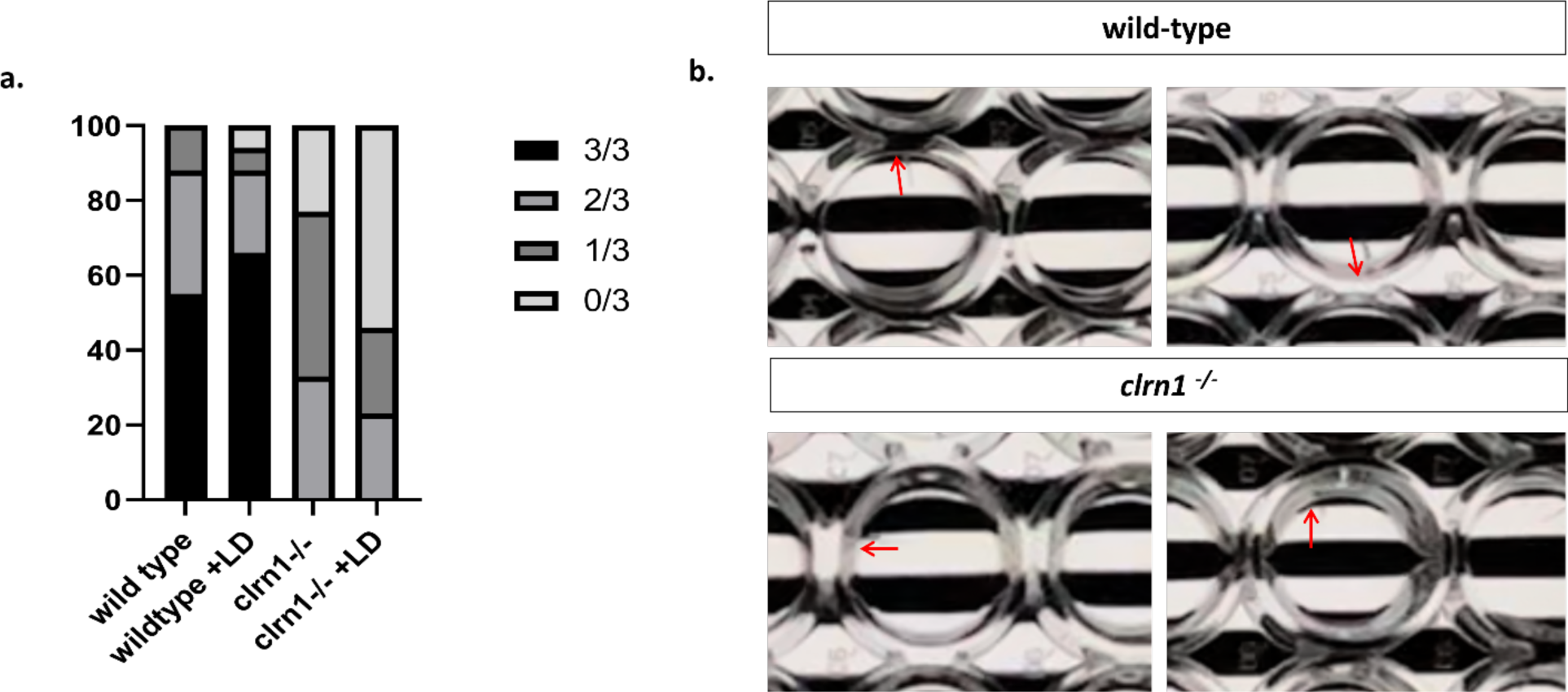
OMR responses in wild-type and *clrn1^-/-^* zebrafish maintained in standard or high-intensity light conditions 5 to 7 dpf. Distribution of OMR responses in wild-type and *clrn1^-/-^* control or light damaged zebrafish (G) Representative examples of wild-type and *clrn1^-/-^* responses following stimulus direction change (H).

**Supplemental Figure 7:**
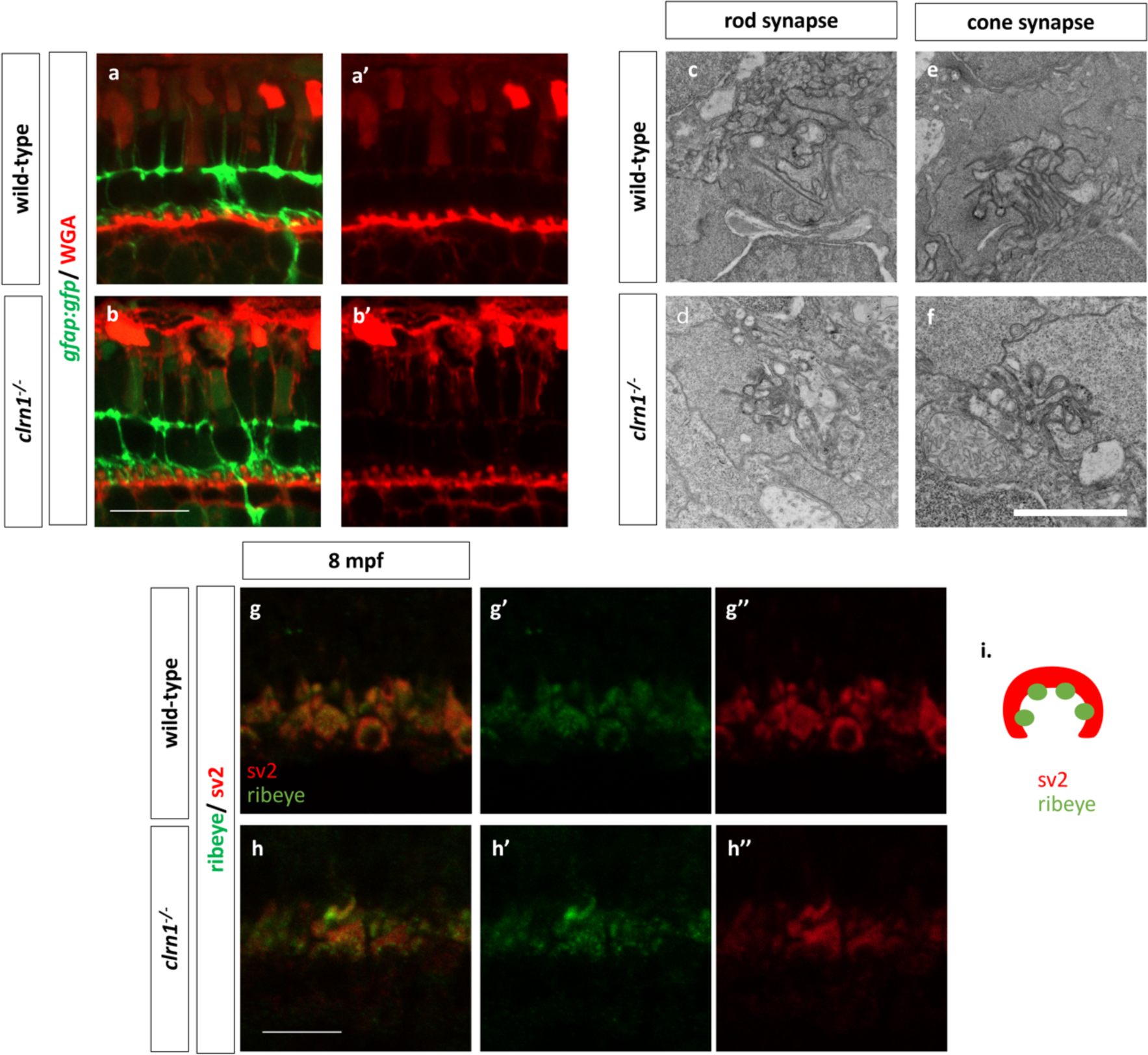
Clrn1 is not required for synaptic development. (a, b) WGA (red) staining marks the lectin rich membrane of pre- and post-synaptic terminals, while Müller glia expressed GFP highlight cellular association with each synapse. There is no discernible difference between wild-type (a) and *clrn1^-/-^* zebrafish (b) at 7dpf. (c-f) Electron micrographs of rod and cone synapses in 7 dpf larvae. There were no discernable differences between wild-type rod (c) and cone (e) synapses compared to *clrn1^-/-^* rod (d) and cone (f) synapses at 7 dpf. (g-h) Anti-SV2 (red) and anti–ribeye (green) marks photoreceptor synapses. Analysis of protein localization and structure revealed no defects in synaptic vesicle localization and ribbon synapse formation in 8 mpf wild-type (g) and *clrn1^-/-^*zebrafish (h). (a-b, g-h) Scale bars: 10 µm, 1 µm (c-f), (Figure a-b, n=10 per group; Figure c-d n=3 per group, Figure g-h n=6-8 per group)

**Supplemental Figure 8:**
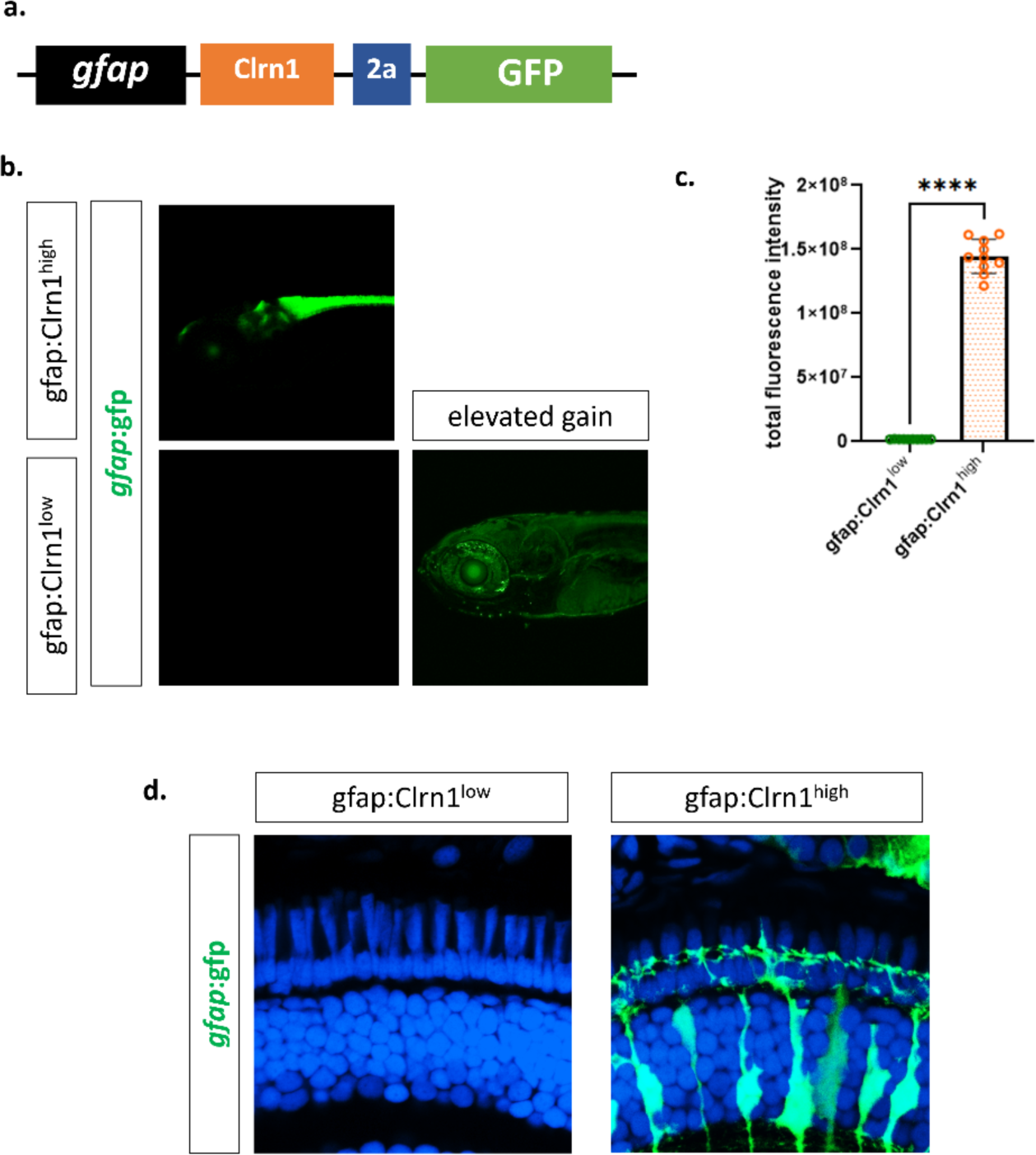
Expression level difference between Clrn1 high and low re-expression in Müller glia transgenic lines. Representation of MG specific Clrn1 expression transgene (a) Representative image of the *Tg(gfap:Clrn1^low^)* and *Tg(gfap:Clrn1^high^)* and Representative image of *Tg(gfap:Clrn1^low^)* at a higher gain to show reporter expression (b). Quantification of total GFP fluorescence intensity in the *Tg(gfap:Clrn1^high^)* and *Tg(gfap:Clrn1^low^)* (c). Transverse sections of 7dpf *Tg(gfap:Clrn1^low^* and *Tg(gfap:Clrn1^high^)* highlighting the loss of GFP signal in *Tg(gfap:Clrn1^low^)* zebrafish tissue post fixation and processing (****p<0.001; Unpaired Students T-test). (n=10)

**Supplemental Figure 9:**
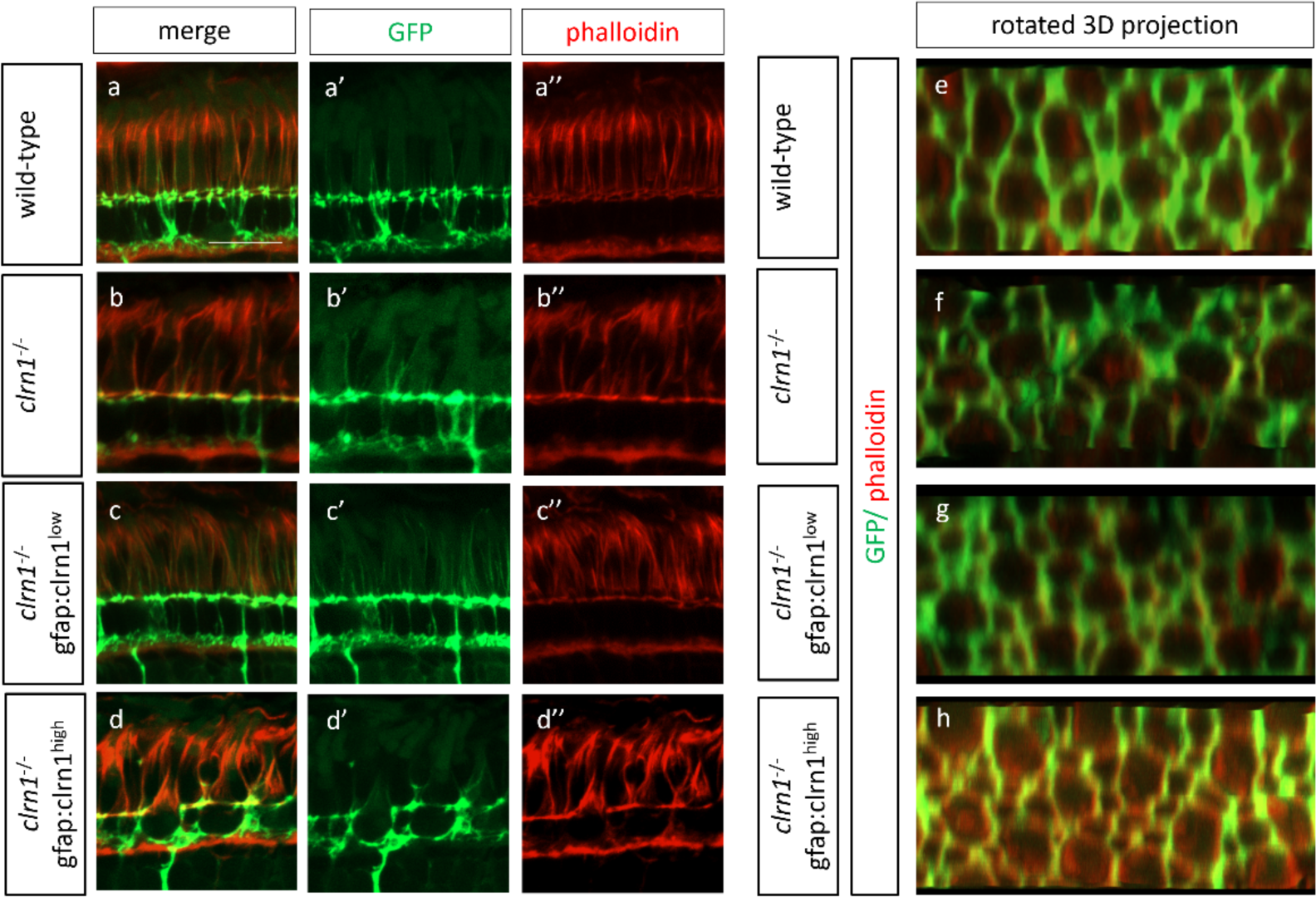
Expression level of clrn1 in Müller glia affects the structure of Müller glia at the OLM and projections of apical microvilli. Comparison of Müller glia structure and actin staining along photoreceptor outersegments (a-d) and at the OLM (e-h) in wild-type (a, e), *clrn1^-/-^* (b, f), *clrn1^-/-^ Tg(gfap:Clrn1^low^)* (c, g), and *clrn1^-/-^ Tg(gfap:Clrn1^low^)* (d, f). To create images of the OLM, 3D projections from Z-stacks were generated and rotated in ImageJ. Comparison of Müller glia structure reveals that re-expression Clrn1 in *Tg(gfap:Clrn1^low^) clrn1^-/-^* zebrafish corrects apical microvilli projects in Müller glia while higher re-expression significantly alters Müller glia structures and affects actin stain along outersegments. Scale Bar: 10 µm (Figure a-b, n=10-15 per group, Figure c-d n=3 per group)

**Supplemental Figure 10.**
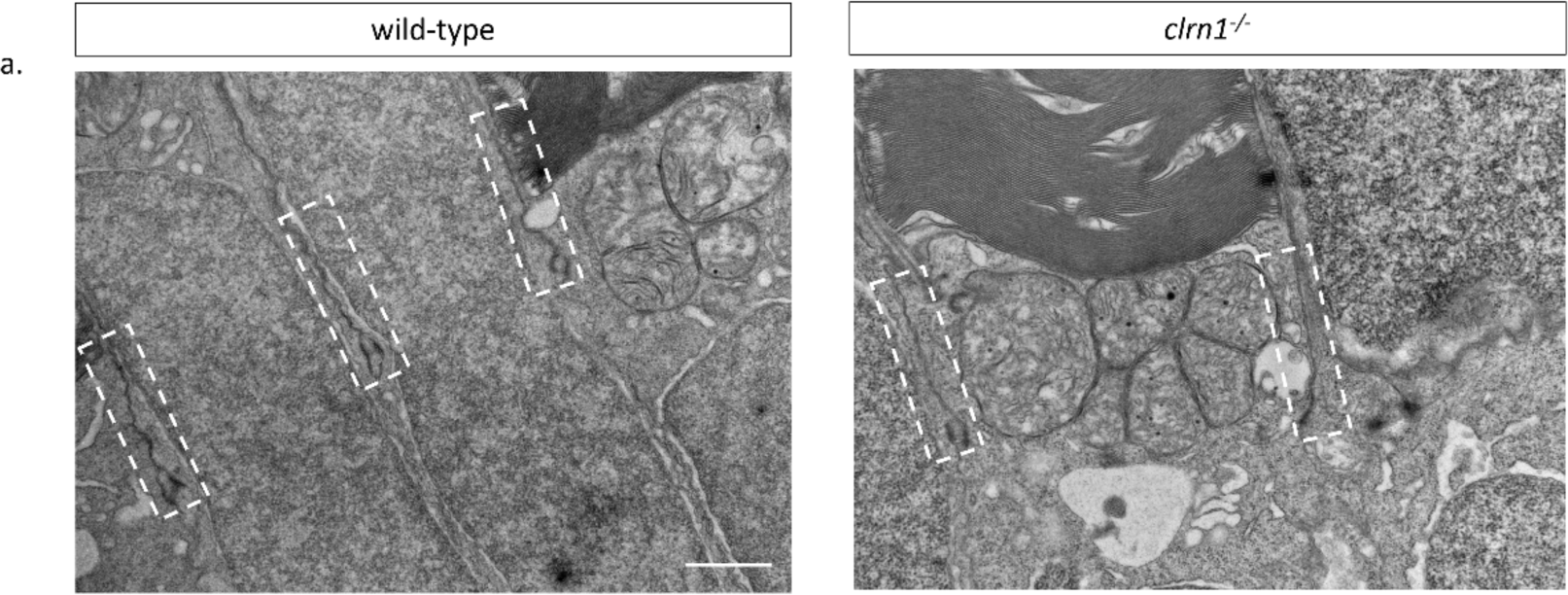
Analysis of photoreceptor junction by TEM. Representative images of the photoreceptor junctions for wild-type and *clrn1^-/-^* at 7dpf. White Boxes highlight junctions in the OLM Scale Bar: 1 uM

**Supplemental Figure 11:**
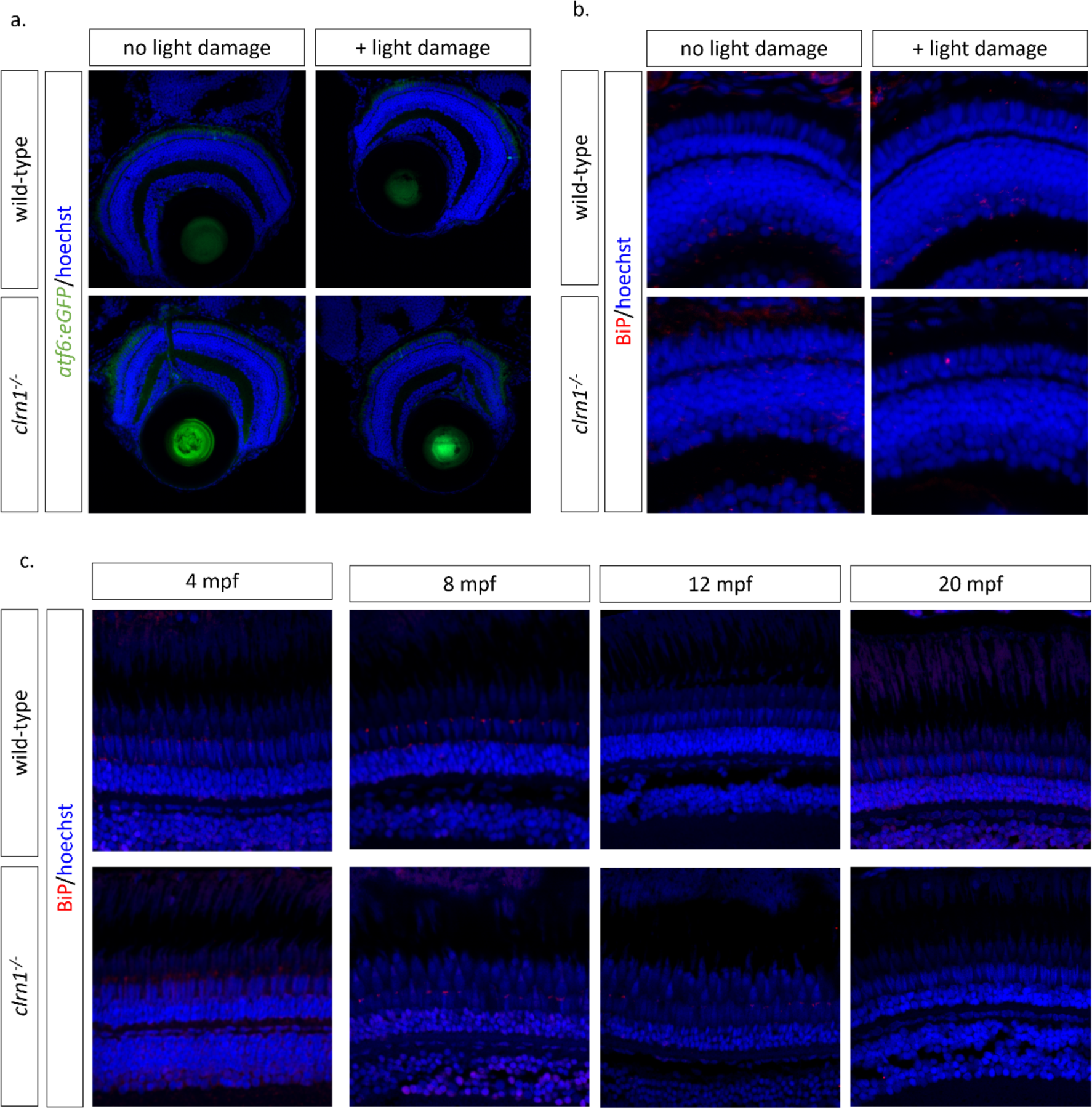
Markers of ER stress pathway activation are not elevated with the loss of Clrn1. Analysis of *Tg(atf6:eGFP)* reporter activation (a) and BiP (b) staining in 6 dpf larvae with or without 24 hours of light damage on transverse cryosections (a). BiP staining on 4-, 8-, 12-, and 20-mpf wild-type and *clrn1^-/-^*zebrafish (c.)

